# Macrophage-intrinsic MDA5-IRF5 axis drives HIV-1 icRNA-induced inflammatory responses

**DOI:** 10.1101/2024.09.06.611547

**Authors:** Sita Ramaswamy, Hisashi Akiyama, Jacob Berrigan, Andrés Quiñones, Alex Olson, Yunhan Chen, Yan Mei Liang, Andrew J. Henderson, Archana Asundi, Manish Sagar, Suryaram Gummuluru

## Abstract

Despite effective antiretroviral therapy (ART), transcriptionally competent HIV-1 reservoirs persist and contribute to persistent immune activation in people living with HIV (PWH). HIV-1-infected macrophages are important mediators of chronic innate immune activation, though mechanisms remain unclear. We previously reported that nuclear export and cytoplasmic expression of HIV-1 intron-containing RNA (icRNA) activates mitochondrial antiviral signaling protein (MAVS)-mediated type I interferon (IFN) responses in macrophages. In this study, we demonstrate an essential role of melanoma differentiation-associated protein 5 (MDA5) in sensing HIV-1 icRNA and promoting MAVS-dependent IRF5 activation in macrophages. Suppression of MDA5, but not RIG-I expression nor disruption of endosomal TLR pathway, abrogated HIV-1 icRNA-induced type I IFN responses and IP-10 expression in macrophages. Furthermore, induction of IP-10 in macrophages upon HIV-1 icRNA sensing by MDA5 was uniquely dependent on IRF5. Additionally, monocytes and MDMs from older (>50 years) individuals exhibit constitutively higher levels of IRF5 expression compared to younger (<35 years) individuals, and HIV-1 icRNA induced IP-10 expression was significantly enhanced in older macrophages, which was attenuated upon ablation of IRF5 expression suggesting that IRF5 functions as a major mediator of pro-inflammatory response downstream of MDA5-dependent HIV-1 icRNA sensing, dysregulation of which might contribute to chronic inflammation in older PWH.

## Introduction

HIV-1 remains a global burden with approximately 39 million people living with HIV (PWH) as of 2022 (UNAIDS/WHO, 2023). Although antiretroviral therapy (ART) has been successful in suppressing virus replication to undetectable levels and extending the lifespan of PWH, systemic inflammatory responses in ART-suppressed PWH remain elevated (1–3). Additionally, chronic systemic inflammation increases the risk for diseases associated with aging such as neurocognitive disorders, cancer, and coronary artery disease (4–7) which account for most of the morbidity and mortality in virologically-suppressed PWH (8) and 5 to 10 years loss in life expectancy compared to risk-adjusted people without HIV (9).

The exact mechanism by which chronic HIV-1 infection contributes to systemic inflammation in virologically-suppressed PWH is unclear. It is well established that early in the course of infection, viral reservoirs are established in long-lived cell populations such as memory CD4+ T cells and macrophages (10, 11). Tissue-resident macrophages located in peripheral lymphoid tissues, liver, brain, lungs, and mucosal tissues harbor HIV RNA and DNA (12–19), and remain persistently infected even during suppressive ART (19–21). While ART reduces peripheral viremia, currently available ART regimens do not suppress viral transcription, and a subset of cells containing integrated provirus remain transcriptionally competent (22). As a result, both viral RNAs and proteins have also been detected in lymph nodes and CNS of ART-suppressed patients (23–25) and are hypothesized to act as pathogen-associated molecular patterns (PAMPs) that can induce persistent inflammatory responses. In concordance with this hypothesis, several recent studies have utilized patient cohort samples to show correlations of HIV-1 RNA/DNA persistence with elevated innate immune markers, and chronic inflammation (26–28).

Chronic immune activation as a consequence of persistent HIV-1 infection is compounded by the low-grade chronic inflammation that occurs during aging, and a combination of these phenotypes is referred to as “HIV inflammaging” (29, 30). This innate immune aging phenotype is proposed to be mediated by cells in the tissue and stroma responding to diverse stimuli within the tissue microenvironment and at the systemic level (31, 32). Tissue-resident macrophages are potentially the primary sensors of cellular injury or infection and responsible for elevated secretion of infection-or injury-induced inflammatory cytokines such as IL-6, sCD14 and sCD163 (33). In addition, aged macrophages have diminished phagocytic capabilities, resulting in a failure to resolve inflammation (34, 35), and have been primarily associated with tissue pathology and age-associated end-organ diseases (36–39). Several studies have indicated that the population of monocytes and macrophages increases with age in many tissue compartments and the polarization of these cells towards an inflammatory phenotype is also enhanced in older people without HIV (40–42) and in older PWH (43). How HIV infection contributes to premature and accelerated aging of the innate immune system has remained unclear.

Previous findings from our group and others showed that cytoplasmic expression of HIV-1 intron-containing RNA (icRNA) triggers induction of interferon stimulated gene (ISG) expression and type I interferons in macrophages and dendritic cells (44, 45). Nuclear export of HIV-1 icRNA is dependent on HIV-1 protein Rev which recognizes and binds a structured domain of HIV-1 RNA referred to as the Rev Response Element (RRE) (46). Upon Rev binding to RRE, a nuclear export factor, CRM1, is recruited to Rev and the Rev/RRE complex is shuttled out of the nucleus into the cytosol (47). Sensing of HIV-1 icRNA in the cytoplasm results in the activation of the mitochondrial antiviral signaling protein, MAVS, initiating a downstream signaling response and innate immune activation (44, 45). However, the sensor involved in HIV-1 icRNA sensing in macrophages remains unknown.

Retinoic acid-inducible gene I (RIG-I) like receptors (RLRs) are cytosolic sensors that detect double stranded RNAs (dsRNA) and mediate an interferon response to viral infections (48). RLRs include RIG-I, melanoma differentiation-associated protein 5 (MDA5), and laboratory of genetics and physiology 2 (LGP-2), though only RIG-I and MDA5 have been shown to sense dsRNAs in a sequence-independent manner (49, 50), Oligomerization of MAVS allows for the recruitment of various E3 ubiquitin ligases, TNF receptor associated factors (TRAFs) and cytosolic kinases, which in turn leads to activation of transcription factors such as NF-κB and interferon regulatory factors (IRFs) (51). In this study, we highlight a critical role of MDA5 as a sensor of HIV-1 icRNA in human macrophages and IRF5 in mediating MDA5/MAVS-dependent type I IFN responses and pro-inflammatory cytokine production. Interestingly, monocytes and MDMs from older donors (ages 50 and above, >50 yo) displayed elevated levels of constitutive IRF5 expression compared to younger donors (ages 35 and younger, <35 yo), which correlated with higher levels of HIV-1 icRNA-induced IP-10 production in MDMs. Taken together, these findings offer mechanistic insights into the understanding of how persistently-expressed HIV-1 icRNA can promote chronic expression of proinflammatory cytokines, and might contribute to inflammaging in older PWH.

## Results

### Rev/CRM1-dependent nuclear export and MAVS are required for HIV-1 icRNA sensing in THP-1/PMA macrophages

Previous work from our group and others identified that Rev/CRM1-mediated export of HIV-1 icRNA in MDMs and DCs was required for induction of MAVS-dependent innate immune responses, though the identity of the RNA sensor or the signaling pathway downstream of MAVS activation has remained unclear (44, 45). To further characterize this signaling pathway in a tractable system, we employed monocytic THP-1 cells and differentiated them with phorbol 12-myristate-13-acetate (PMA) to macrophage-like cells (52). THP-1/PMA macrophages were infected with VSV-G-pseudotyped single-cycle HIV-1 encoding GFP as a reporter in place of nef (LaiΔenvGFP/G) (Fig. 1A), and pro-inflammatory cytokine, CXCL10 (IP-10) secretion in infected cultures was employed as a quantitative measure of infection-induced innate immune activation. Similar to findings in MDMs, establishment of productive infection in THP-1/PMA macrophages resulted in robust secretion of IP-10 (Fig. 1B), which was abrogated in the presence of inhibitors that target reverse transcriptase (efavirenz), integrase (raltegravir), or viral transcription (spironolactone) (53–55) (Fig. 1B). Furthermore, infection with HIV-1/M10 mutant (Fig. 1C) that is incapable of exporting HIV-1 icRNA via the Rev/CRM1 pathway(46), or infection in the presence of CRM1 inhibitor (KPT330) also failed to induce IP-10 secretion (Fig. 1B and D). Knock-down of MAVS expression in THP-1/PMA macrophages (Fig. 1E, Sup. Fig. 1A-E) significantly decreased IP-10 secretion in virus-infected cells (Fig. 1F-G). Collectively, these findings suggest that sensing of a late post-transcriptional step of the HIV-1 replication cycle induces innate immune responses in THP-1/PMA macrophages and recapitulate previous findings in HIV-1-infected MDMs and DCs (44, 45).

**Figure 1:**
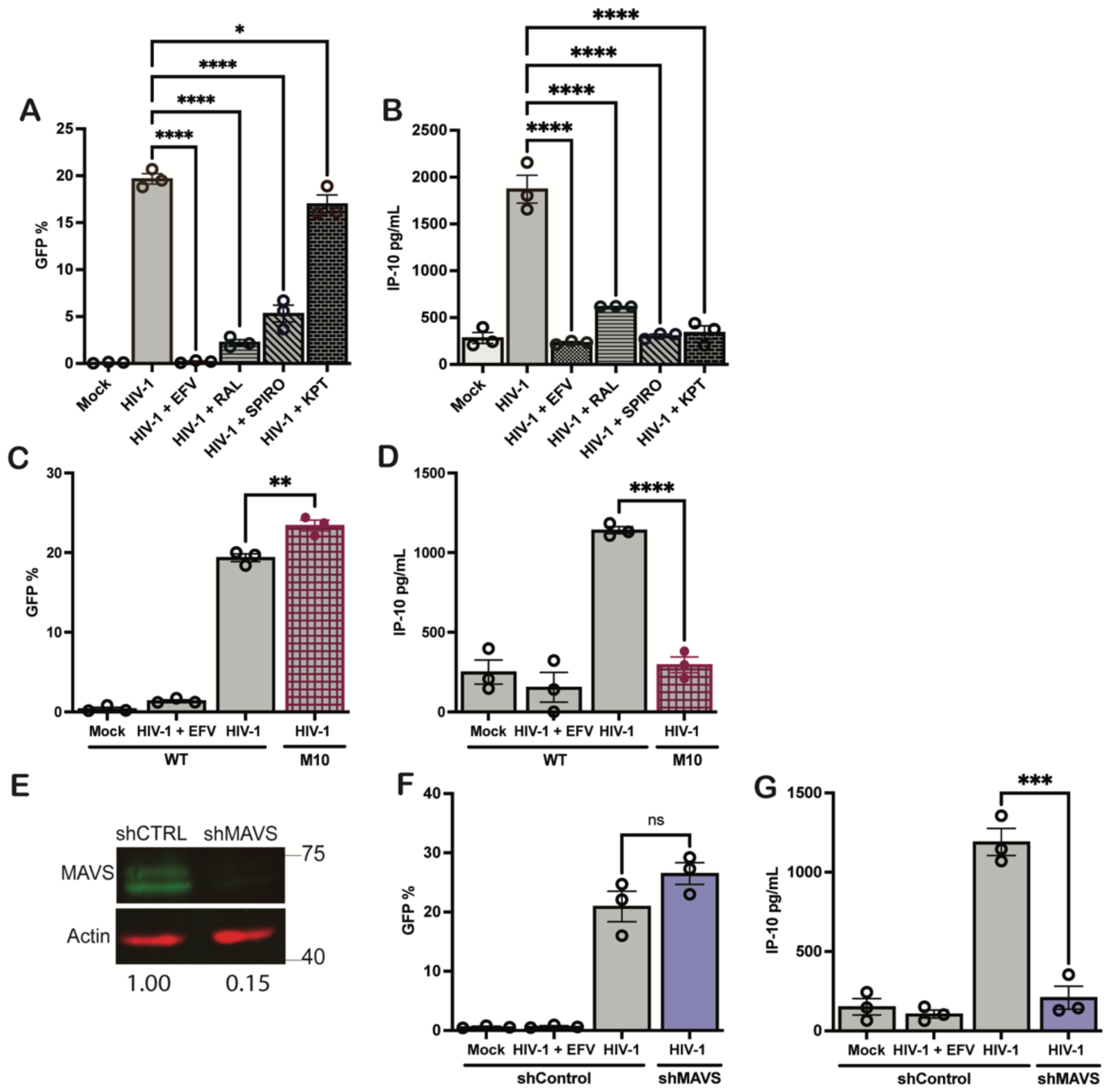
MAYS-mediated sensing of cytoplasmic icRNA triggers an innate immune response. THP-1/PMA macrophages were infected with Lailienv GFP/G (MOI 2) in the presence or absence of efaverinz (EFV, 1 µM), raltegravir (RAL, 30 µM), spironolactone (Spiro, 100 nM), and KPT335 (KPT, **1 µM).** Cells and culture supernatants harvested at 3 dpi for (A) flow cytometry analysis to measure infection levels (¾GFP+) and (B) ELISA to measure IP-10 secretion. Infection levels (C) and IP-10 secretion (D) in THP-1/PMA macrophages infected with either WT (LailienvGFP/G) or HIV-1/M10 were determined at 3 dpi by flow cytometry and ELISA, respectively. (E) MAVS expression in THP1 cells transduced with shCTRL or shMAVS lentivectors was determined by western blot analysis. (F, G) LaiiienvGFP/G infected THP-1/PMA macrophages and cell supernatants were harvested 3 dpi for flow cytometry analysis (F) and ELISA (G) to measure infection establishment (¾GFP+) and IP-10 secretion. Data is displayed as the means ± SEM with each dot representing an independent experiment. Statistical significance assessed via 1-way ANOVA with Dunnet’s multiple comparisons test (A-B), unpaired t-test (C-O, F-G). *: p < 0.05; **: p <0.01, ***: p < 0.001, ****: p < 0.0001, ns = not significant.

### MDA5 is required for HIV-1 induced immune response in macrophages

In mammalian cells, detection of viral RNAs in the cytosol and endosomes is surveilled by a limited number of nucleic acid receptors. In order to identify the nucleic acid sensing mechanism required for detection of HIV-1 icRNA, we generated THP-1 cell lines with stable knock-down of RIG-I (sensor of short, blunt ended dsRNAs with uncapped 5’ triphosphate group), MDA5 (sensor of long dsRNAs), or UNC93B1 (chaperone protein required for endosomal TLR 3/7/8 function) expression in THP-1 cells via lentiviral transduction of shRNA (56). Knockdown efficiency was measured at the mRNA level via RT-qPCR (Fig 2A), while decrease in RLR or endosomal TLR activity was functionally validated by measuring diminished IP-10 secretion in knockdown cells in response to synthetic or viral RLR or TLR agonists (Sup Fig. S1F-J). While single-cycle HIV-1 (Lai∆envGFP/G) infection of THP-1/PMA macrophages with reduced expression of RIG-I, MDA5 or UNC93B1 was unaffected compared to control shRNA expressing cells (Fig. 2B), we found that only MDA5, but not RIG-1 or UNC93B1, knockdown resulted in significant downregulation of HIV-1 icRNA-induced IP-10 production (Fig. 2C). We next recapitulated these findings in primary MDMs. MDMs were transfected with pooled siRNAs against RIG-I, MDA5 or UNC93B1 and reduction in mRNA expression of RLRs and TLR chaperone was verified via RT-qPCR (Fig. 2D). Upon infection of MDMs with Lai∆envGFP/G, we observed downregulation of IP-10 and IFN∆ mRNA expression upon MDA5 knockdown (Fig. 2E-F, Sup Fig. 1K), but not upon knockdown of either RIG-I or UNC93B1 (Fig. 2F & Sup. Fig. 1K). Taken together, these results suggest that MDA5 is required for induction of innate immune response upon cytoplasmic HIV-1 icRNA expression in MDMs.

**Figure 2:**
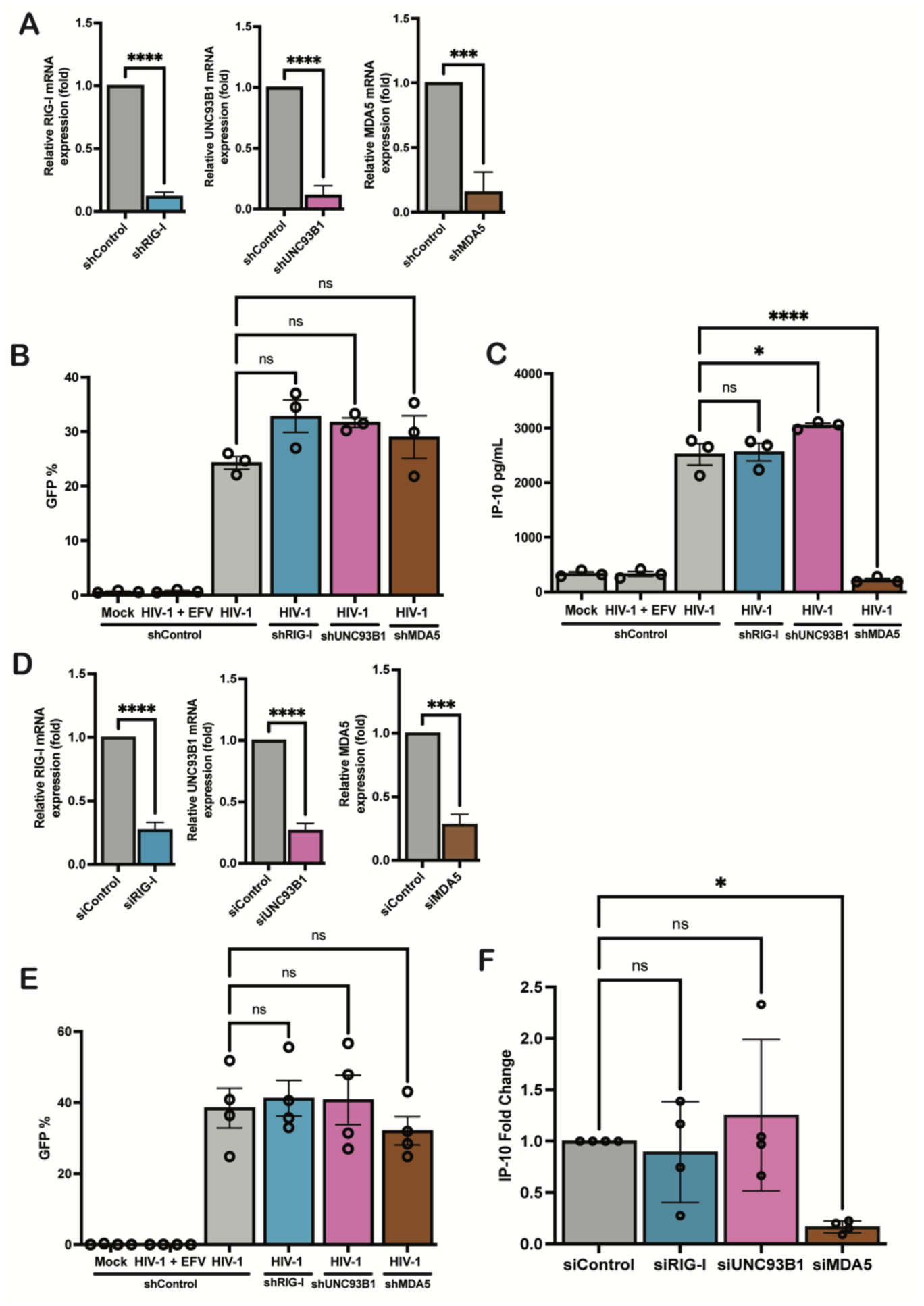
MDA5 is required for HIV-1 induced innate immune response in macrophages. (A) RIG-I, UNC93B1, and MDA5 expression in THP-1 cells transduced with RIG-I, UNC93B1, MDA5 or Ctrl shRNA-expressing lentivectors was quantified via RT-qPCR. (B) THP-1/PMA knockdown cell lines were infected with Lai envGF­ P/G at MOI 2 and harvested 3 dpi for infection establishment (¾GFP+) via flow cytometry. (C) Supernatants from infected THP-1/PMA cells were used to assess IP-10 secretion via ELISA. (D) MDMs were transfected with siRNA targeting RIG-I, UNC93B1, and MDA5 for 2 days and knockdown of RIG-I, UNC93B1, and MDA5 expression was assessed via RT-qPCR. MDMs were infected LaiMnvGFP/G at MOI 1 in the presence of dNs and harvested at 2 dpi for analysis of infection efficiency via flow cytometry (E) and IP-10 mRNA expression via RT-qPCR (F). Data is displayed as the means ± SEM with each dot representing an independent experiment (A-C) or cells from an independent donor (D-F). Statistical significance assessed via unpaired t-test (A, D), 1-way ANOVA with Dunnett’s multiple comparisons (B-C, E-F). *: p < 0.05; ***: p < 0.001, ****: p < 0.0001, ns = not significant.

### MDA5 recognizes unspliced HIV-1 RNA

In order to investigate the interaction of MDA5 with HIV-1 icRNA, HEK293T cells were infected with HIV-1 (Laiι1env GFP/G) in the presence or absence of efaverinz and then transfected with plasmids expressing either Flag-epitope tagged MDA5 or RIG-I. To ensure equivalent expression level of MDA5 and RIG-I, HEK293T cells were transfected with varying amounts of expression plasmids, and transfection conditions that resulted in similar levels of RIG-I or MDA5 expression were utilized for RNA immunoprecipitation analysis (Sup. Fig 1L). RNA-protein interactions were stabilized in situ via UV crosslinking prior to fractionation of cell lysates to nuclear and cytoplasmic fractions, and immunoprecipitation of ribonucleoprotein particles (RNPs). Immunoblots confirmed equivalent MDA5 and RIG-I expression in cytoplasmic fractions (Fig. 3A), and immunoprecipitation by anti-flag mAb (Fig. 3B). Immunopreciptated RNA was quantified via RT-qPCR with primers complementary to *gag* (unspliced RNA, usRNA), *tat-rev* (multiply spliced RNA, msRNA), *GAPDH* or *actin.* In concordance with the robust functional attenuation of HIV-1 icRNA sensing upon knock-down of MDA5 expression (Fig. 2), HIV-1 usRNA was significantly enriched in RNPs immunoprecipitated with anti-flag mAb in MDA5-flag expressing cells (mean ∼20-fold enrichment for MDA5), but not in RIG-I flag expressing cells or control IgG immunopreciptates (Fig. 3B). In contrast, no significant differences were observed in the level of msRNA co-immunopreciptated with MDA5 or RIG-1. Specificity of HIV-1 usRNA immunopreciptation by MDA5 was confirmed by the absence of GAPDH or actin RNAs in anti-flag or IgG immunoprecipitates (Fig. 3B). These findings are in agreement with recently published studies that demonstrate the ability of MDA5 to specifically recognize HIV-1 usRNA in virus-infected dendritic cells (57, 58). Taken together, these results suggest that MDA5 specifically interacts with HIV-1 icRNA and that MDA5 sensing of HIV-1 icRNA triggers innate immune activation in macrophages.

**Figure 3:**
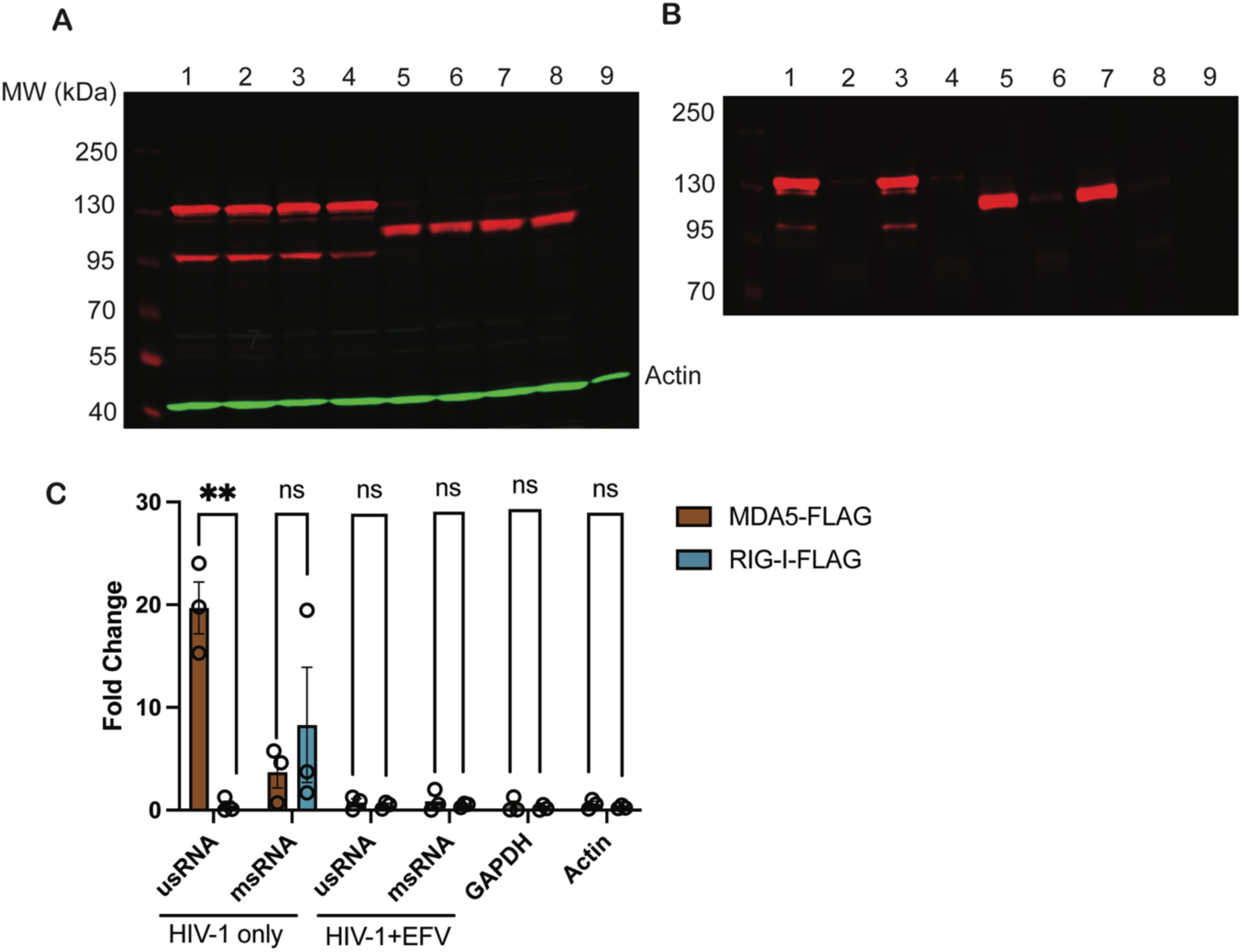
MDA5 recognizes unspliced HIV-1 RNA. HEK293T cells were infected with LaiMnvGFP/G at MOI 1 and transfected with either MDA5-Flag or RIG-I-Flag at 24 h post infection. Cytoplasmic fractions were immunoprecipitated with either anti-Flag mAb or lgG coated beads.(A) Input and (B) IP lysates were run on western blot to ensure equivalent levels of transfection and immunoprecipitation among conditions (1) HIV MDA5-Flag (2) HIV MDA5-lgG (3) HIV+EFV MDA5-Flag (4) HIV+EFV MDA5-lgG (5) HIV RIG-I-Flag (6) HIV RIG-I-lgG (7) HIV+EFV RIG-I-Flag (8) HIV+EFV RIG-I-lgG (9) Mock (C) RT-qPCR analysis for HIV-1 usRNA, HIV-1 msRNA, GAPDH or Actin mRNA in transfected and infected 293Ts in the presence or absence of efaverinz. Fold enrichment of immunoprecipitated RNA reported as RNA transcripts detected from each IP condition using anti-flag mAb or control lgG to the input amount. Data is displayed as means ± SEM with each dot representing a different experiment. Statistical significance assessed via unpaired !-tests (C). **: p < 0.01, ns = not significant.

### IRF5 is a mediator of HIV-1 induced IP-10 production in macrophages

Upon sensing of viral RNAs, CARD-dependent interactions of MDA5 with MAVS (59, 60) leads to MAVS oligomerization and activation of downstream transcription factors including NF-κB and IRFs(60). Within the IRF family of transcription factors, IRF3, 5, and 7 have well-described roles in mediating antiviral and inflammatory responses downstream of diverse viral infections (61). To characterize the roles of these IRFs in mediating HIV-1 icRNA-induced innate immune response in macrophages, we generated THP-1 cells with stable knockdown of IRF3, IRF5 or IRF7 expression using lentiviral shRNA transduction. We confirmed knockdown of these IRFs via Western blot (Fig 4A) and RT-qPCR (Sup. Fig 2A-C), and observed similar efficiency of decrease in IRF3, IRF5 and IRF7 expression. Functional knockdown was characterized by measuring response to 3p-hpRNA (RIG-I agonist) or LPS (TLR4 agonist) treatment. Decreased IRF3, 5 or 7 expression in THP-1/PMA macrophages led to attenuated responses to both 3p-hpRNA and LPS (Sup. Fig 2D-E). THP-1/PMA macrophages were infected with Lai∆envGFP/G, and infection efficiency was assessed via flow cytometry 3 days post infection. We found there was no significant effect of IRF3, IRF5, or IRF7 knockdown on HIV-1 infection (Fig. 4B). Culture supernatants were harvested 3 days post infection and analyzed for IP-10 production via ELISA. While knock down of IRF5 expression resulted in robust downregulation of HIV-1 icRNA-induced IP-10 production, decrease in IRF3 or IRF7 expression led to a modest though significant attenuation of IP-10 secretion (Fig. 4C). Interestingly, requirement of IRFs for IP-10 secretion was dependent on the nature of viral PAMPs and the RLR sensing pathway, as IRF3, but not IRF5 or IRF7, was selectively required for Sendai virus-induced RIG-I-dependent IP-10 secretion (Sup. Fig.2F-G). To confirm these findings in primary MDMs, expression of IRF3, IRF5 and IRF7 was knocked-down via transient transfection with pooled siRNAs (Fig. 4D, Sup Fig. 2H-J) prior to infection with Lai∆envGFP/G. While knock-down of IRF3, IRF5 or IRF7 expression had no impact on the efficiency of infection establishment (Fig. 4E), we observed significant downregulation in IP-10 expression upon IRF5 or IRF3 knockdown, but not IRF7, in HIV-infected MDMs (Fig. 4F). In contrast to IP-10, expression of IFNβ was robustly attenuated by knockdown of IRF3, IRF5 or IRF7 expression in HIV-1-infected MDMs (Sup Fig. 2K), suggesting a selective and non-redundant role of IRF5 in mediating a proinflammatory response to HIV-1 infection in macrophages.

**Figure 4:**
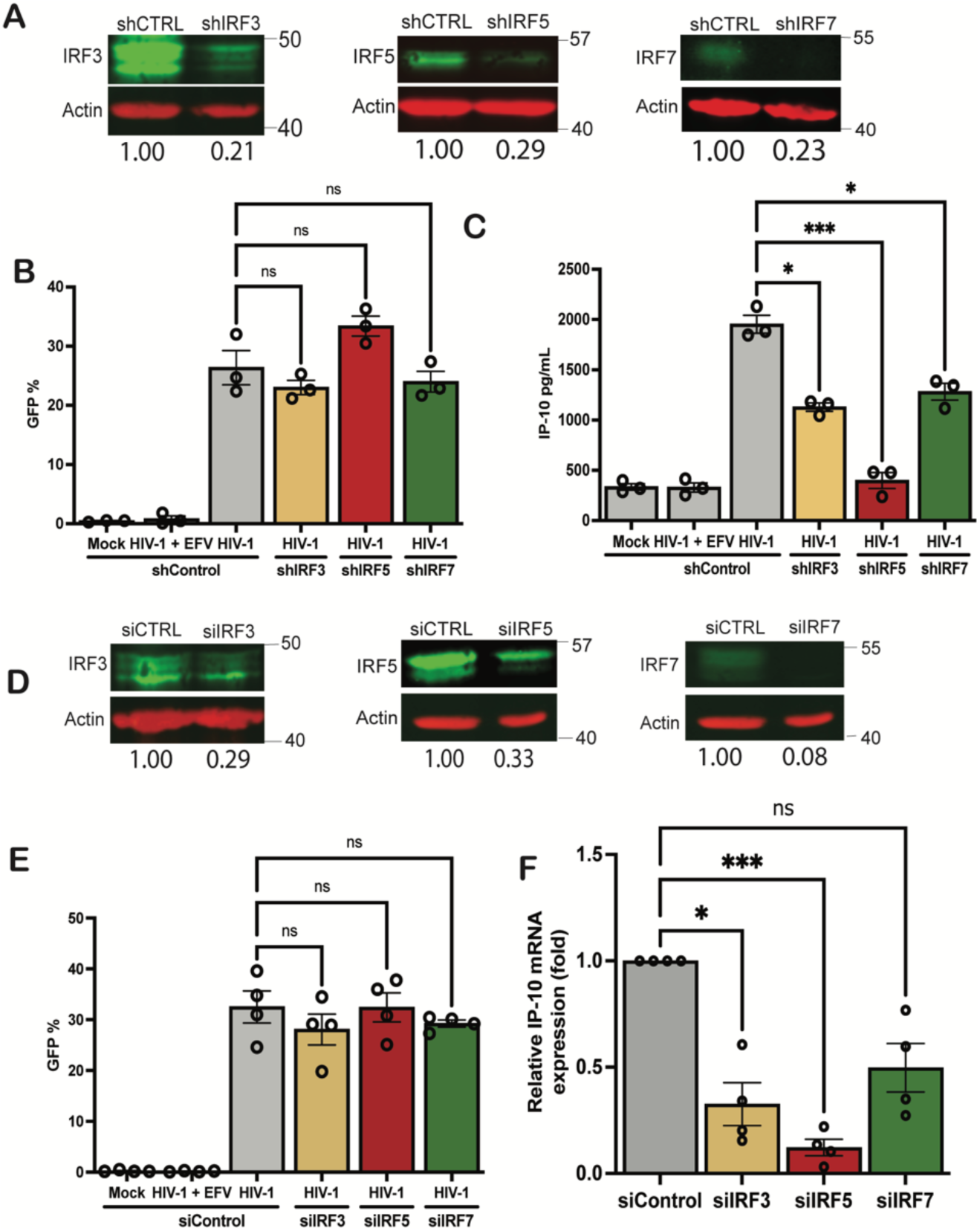
IRF5 is necessary for HIV-1 icRNA-induced IP-10 expression in macrophages. (A) IRF3, IRF5 and IRF7 expression in THP-1 cells transduced with IRF3, 5, 7 or Ctrl shRNA lentivectors was quantified via western blot analysis and normalized to shControl transduced cells. (B) THP1/PMA macrophages were infected with Lait. envGFP/G at MOI 2 and cells and supernatants harvested 3 dpi for analysis via flow cytometry to assess infection levels (B) and ELISA for IP-10 secretion (C). (D) Expression of IRFs in MDMs transfected with siRNA against IRF3, IRF5, or IRF7 mRNA was assessed via RT-qPCR. (E) MDMs were infected with LaiMnvGFP/G at MOI 1 in the presence of dNs and harvested at 2 dpi for analysis of infection efficiency via flow cytometry (E) and IP-10 expression via RT-qPCR (F). Data is displayed as the means± SEM with each dot representing a different donor. Statistical significance assessed via 1-way ANOVA with Dunnet’s multiple comparisons (B-C, E-F). *: p < 0.05; ***: p < 0.001, ns = not significant.

### TRAF6 and IKK∆ are required for IP-10 production in HIV-1-infected macrophages

Upon sensing of viral RNA by MDA5 and activation of the MAVS signalosome, diverse kinases including IKKβ, IKKε, and TBK1 and ubiquitin ligases such as TRAF2, TRAF5 and TRAF6 are recruited and activated, which in turn post-translationally modify IRFs and NF-κB to drive expression of antiviral genes and inflammatory cytokines (62, 63). These modifications are critical for IRF5 activation (62, 63). It has previously been shown that the E3 ubiquitin ligase, TRAF6, and serine kinase, IKKβ, are required for K63-linked polyubiquitination and phosphorylation events and IRF5 activation downstream of TLR and RLR sensing (64), though their requirement for HIV-1-induced type I IFN responses and IP-10 production has not been assessed. To determine whether IKKβ and TRAF6 were required for IP-10 production upon HIV-1 icRNA sensing in macrophages, we utilized lentiviral shRNA to generate stable TRAF6 or IKKβ knock-down THP-1 cell lines. Knockdown of gene expression of these factors was validated by Western blot and RT-qPCR. (Fig. 5A, Sup Fig. 2L-M). These knockdown cell lines were also validated functionally by measuring IP-10 secretion, which was significantly attenuated in IKKβ or TRAF6 knock-down cells in response to RLR and TLR agonists such as 3p-hpRNA and LPS (Sup Fig. 2N-O). While knockdown of TRAF6 or IKKβ expression resulted in no difference in HIV-1 infection (Fig. 5B), we observed a significant decrease in IP-10 production in HIV-1-infected THP-1/PMA macrophages (Fig. 5C). In contrast, IP-10 secretion was only modestly attenuated in HSV or Sendai virus-infected THP-1/PMA macrophages deficient for TRAF6 or IKKβ expression (Sup Fig. 2P-Q). These results suggest that, in contrast to complementary roles of TRAF2, TRAF5, and TRAF6 downstream of MAVS activation in HSV or Sendai virus-infected cells (43), TRAF6 has a non-redundant role in the induction of IP-10 expression in response to HIV-1 icRNA sensing by MDA5. We next sought to determine the roles of TRAF6 and IKKβ in transducing signals downstream of HIV-1 icRNA sensing in primary MDMs. MDMs were transfected with pooled siRNAs against TRAF6 or IKKβ prior to infection with LaiΔenvGFP/G. Quantitative RT-qPCR and western blot analysis confirmed robust knock-down efficiency of both TRAF6 and IKKβ in MDMs (Fig. 5D, Sup Fig. 2R-S). Similar to the findings in THP-1/PMA macrophages, while knockdown of TRAF6 or IKKβ expression did not attenuate HIV infection (Fig. 5E), there was a robust reduction in HIV-1 icRNA-induced IP-10 and IFNβ mRNA expression upon knockdown of TRAF6 or IKKβ expression in MDMs (Fig. 5F, Sup Fig. 2T), indicating an important role of TRAF6 and IKKβ in MDA5/MAVS-dependent activation of IRF5 for induction of IP-10 and type I IFN responses in HIV-1-infected macrophages.

**Figure 5:**
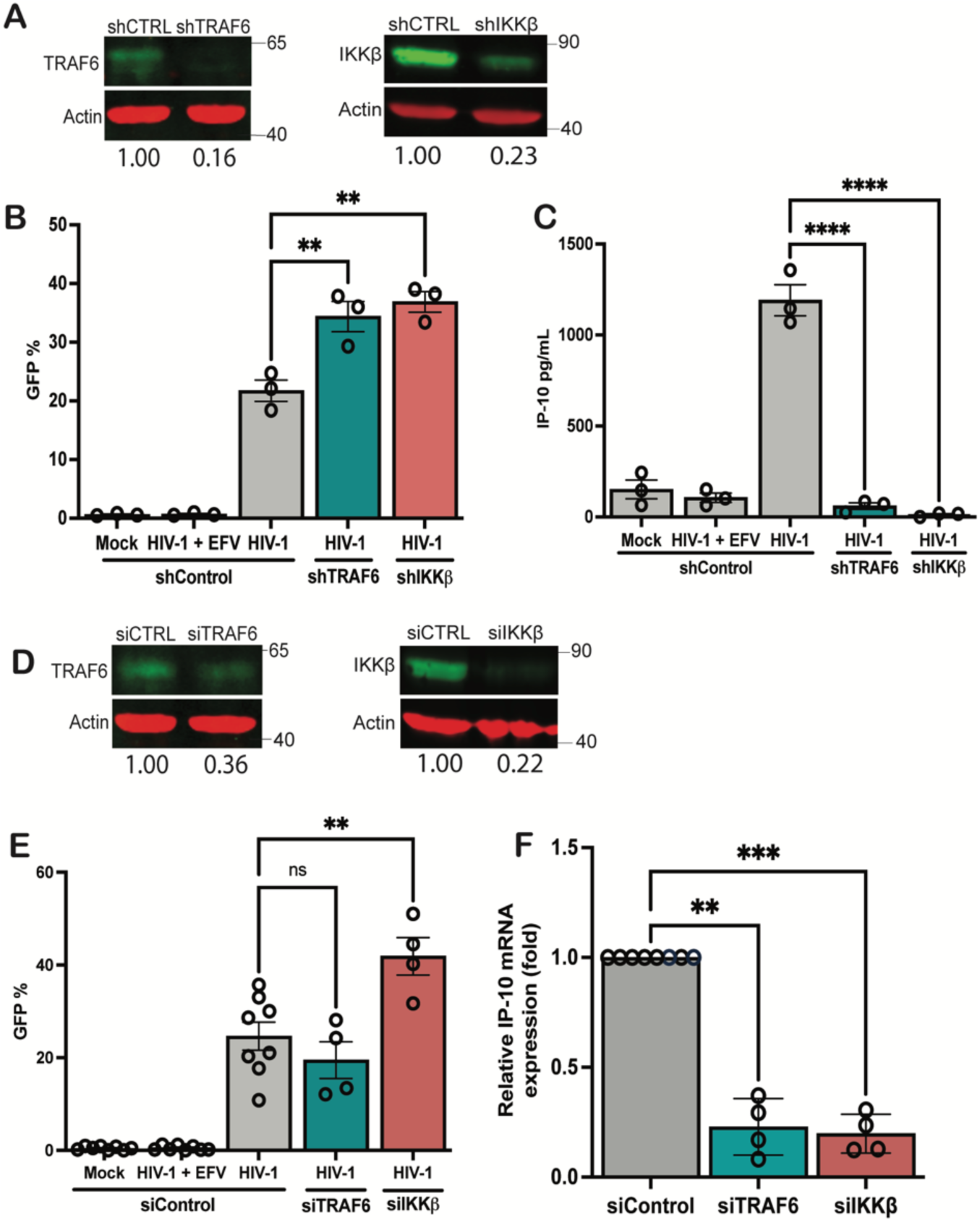
TRAF6 and IKKj3 are required for HIV-1 induced IP-10 expression in macrophages. (A) TRAF6 or IKKj3 expression in THP-1 cells transduced with TRAF6 or IKKb shRNA lentivectors was quantified via western blot analysis and normalized to shControl transduced cells. THP1/PMA macrophages infected with Lai MnvGFP/G at MOI 2 and harvested 3 dpi for analysis via flow cytometry to assess infection levels (B) and ELISA to assess IP-10 secretion (C). (D) MDMs were transfected with siRNA against TRAF6 or IKKj3mRNA for 2 days and knockdown of TRAF6 or IKKj3 was assessed via WB and RT-qPCR. (E) MDMs infected with Lai envGFP/G at MOI 1 in the presence of dNs and harvested at 2 dpi for analysis of infection efficiency via flow cytometry (E) and IP-10 expression by RT-qPCR (F). Data is displayed as the means± SEM with each dot representing a separate experiment (A-C) with cells from independent donors (D-F). Statistical significance assessed via 1-way ANOVA with Dunnet’s multiple comparisons (B-C, E-F). **: p < 0.01, ****: p < 0.0001, ns = not significant.

### HIV-1 infection results in nuclear translocation of IRF5 in THP-1/PMA macrophages

Since IRF5 is an ISG (65), and HIV-1 infection of macrophages induces type I IFN responses, we next sought to determine if IRF5 activation in HIV-infected MDMs occurred directly downstream of MDA5/MAVS sensing of HIV icRNA in a cell-intrinsic manner in infected cells or required activation in bystander uninfected cells. IRF5 exists in an inactive form in the cytosol in unstimulated cells, post-translational modifications of which result in its translocation to the nucleus (66). To visualize IRF5 activation in response to HIV-1 infection in macrophages, we utilized immunofluorescence and confocal microscopy to assess its nuclear localization. THP-1/PMA macrophages were infected with Laiι∆envGFP/G, and stained for intracellular localization of IRF5 expression on coverslips at 3 days post infection. DAPI was utilized as a nuclear stain and HIV infected cells were distinguished by GFP positivity. HIV-1 (WT) infection of THP-1/PMA macrophages resulted in nuclear translocation of IRF5 from cytoplasm in infected cells (Fig. 6A), which was not observed upon infection with HIV-1/M10 mutant (Sup. Fig S4A). In order to quantify nuclear translocation of IRF5, we assessed the mean signal intensity of nuclear IRF5 using CellProfiler (Sup Fig. 3). In HIV-1 (WT)-infected, GFP+ THP-1/PMA macrophages, we observed a significant increase in nuclear IRF5 staining, that was blocked upon pre-treatment with EFV or upon infection with HIV-1/M10 mutant virus (Sup Fig. S4A). Furthermore, nuclear localization of IRF5 was selectively enhanced in GFP^+^ cells with HIV WT infection (Sup Fig. S4B), suggesting that cell-intrinsic HIV-1 icRNA sensing activates IRF5 in infected cells. Importantly, nuclear IRF5 localization in HIV-1 (WT)-infected cells was attenuated upon depletion of MDA5, MAVS, IKK∆, or TRAF6 expression (Fig. 6D-G), but not RIG-I expression (Fig. 6C). These results indicate that MDA5, MAVS, IKK∆, and TRAF6 are required for IRF5 activation and nuclear localization directly downstream of HIV-1 icRNA sensing in macrophages.

**Figure 6:**
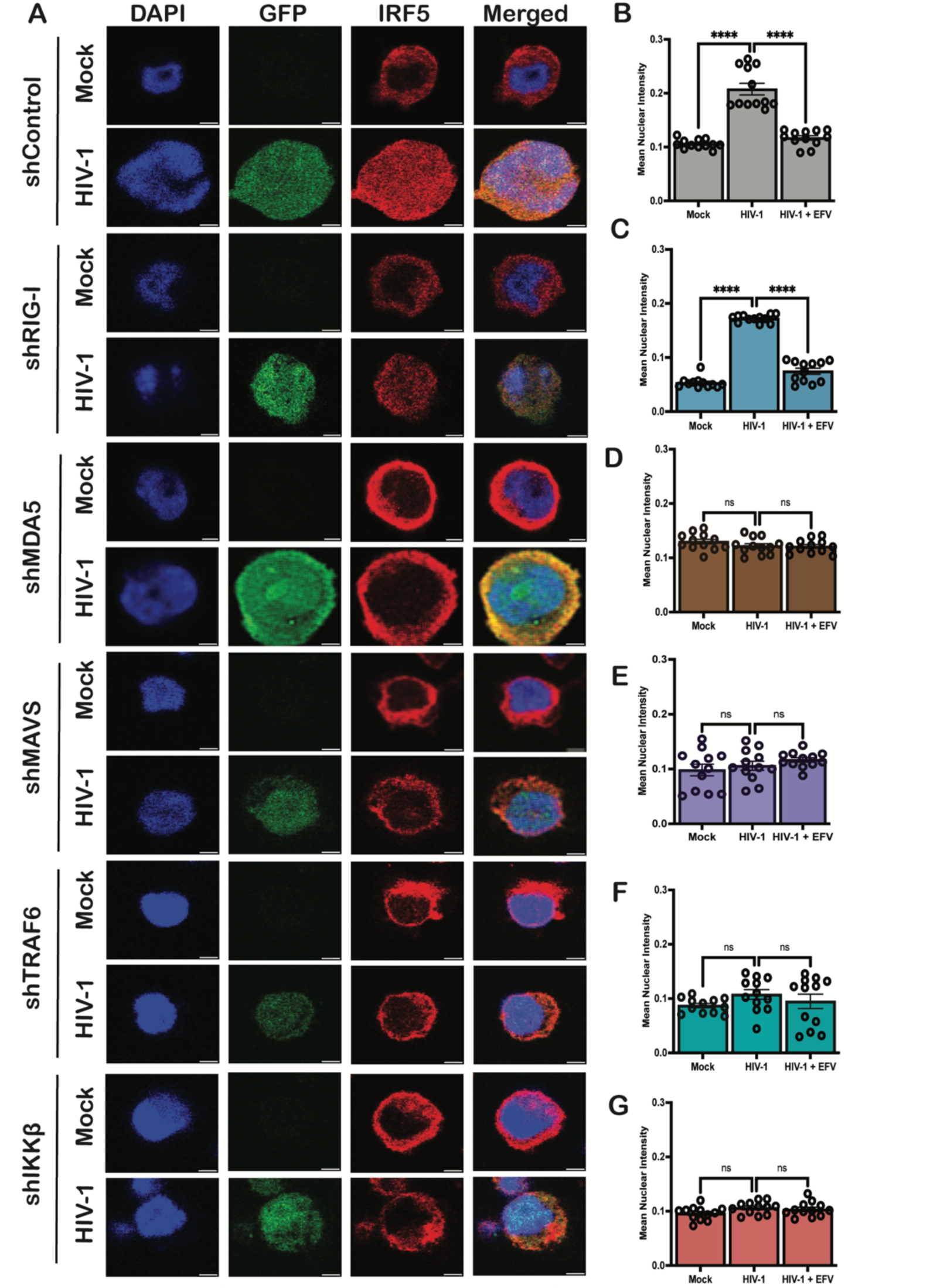
MDA5, MAVS, TRAF6, and IKK13are required for HIV-1 icRNA-induced nuclear localization of IRF5 in macrophages. (A) THP-1/PMA macrophages, and infected with LaiMnvGFP/G at MOI 2 and harvested 3 dpi for analysis via confocal microscopy to assess changes in IRF5 localization. Representative images are shown (scale bar = 5µm). (B-F) Fluorescence microscopy images were quantified via CellProfiler to assess mean pixel intensity of IRF5 staining that colocalized with DAPI (mean nuclear intensity). Images from three independent infection experiments were analyzed and quantified, with each dot representing a field containing approximately 50-150 cells. Statistical significance assessed via Kruskal-Wallis test with Dunn’s multiple comparisons analysis (B-G). ****: p < 0.0001, ns = not significant.

### Macrophages and monocytes isolated from older donors exhibit elevated levels of IRF5

Several cohort studies have demonstrated that older PWH experience co-morbidities and age-associated disease at a greater level compared to age-matched people without HIV (67–69). Despite suppression of viremia, HIV-1 RNA and DNA can persist in long-lived cells such as macrophages and may contribute to the development of co-morbidities in PWH (12–18). We sought to determine whether macrophages isolated from older as compared to younger individuals exhibit a greater inflammatory response to HIV-1 icRNA. To test this hypothesis, we utilized samples from two cohorts, one from the HIV/Aging cohort at Boston Medical Center (BMC) (Sup. Table 1), and the other from NY Biologics Blood Center (Sup. Table 2). Donors were stratified by age as older (>50 years old) or younger (18-35 years old). PBMCs were isolated from whole blood samples from people without HIV, and CD14+ monocytes were collected via positive selection on the day of sample collection, and differentiated to MDMs. In order to investigate basal expression differences in factors involved in sensing of HIV-1 icRNA, we quantified expression levels of MDA5, MAVS, TRAF6, and IRF3/5/7 in CD14+ monocytes and MDMs from younger and older donors. We did not observe any age-associated differences in the expression of MDA5, MAVS, TRAF6, IRF3 or IRF7 expression in peripheral blood CD14+ monocytes and MDMs derived from PBMCs from the NY Biologics Blood Center (Fig. 7A-E). Interestingly, IRF5 expression was constitutively elevated in CD14+ monocytes and MDMs from older individuals (Fig. 7F). We validated this finding by measuring IRF5 protein level via Western blot and found that IRF5 protein expression was significantly elevated in older monocytes and trended to higher expression in older macrophages (Fig. 7G-H).

**Figure 7:**
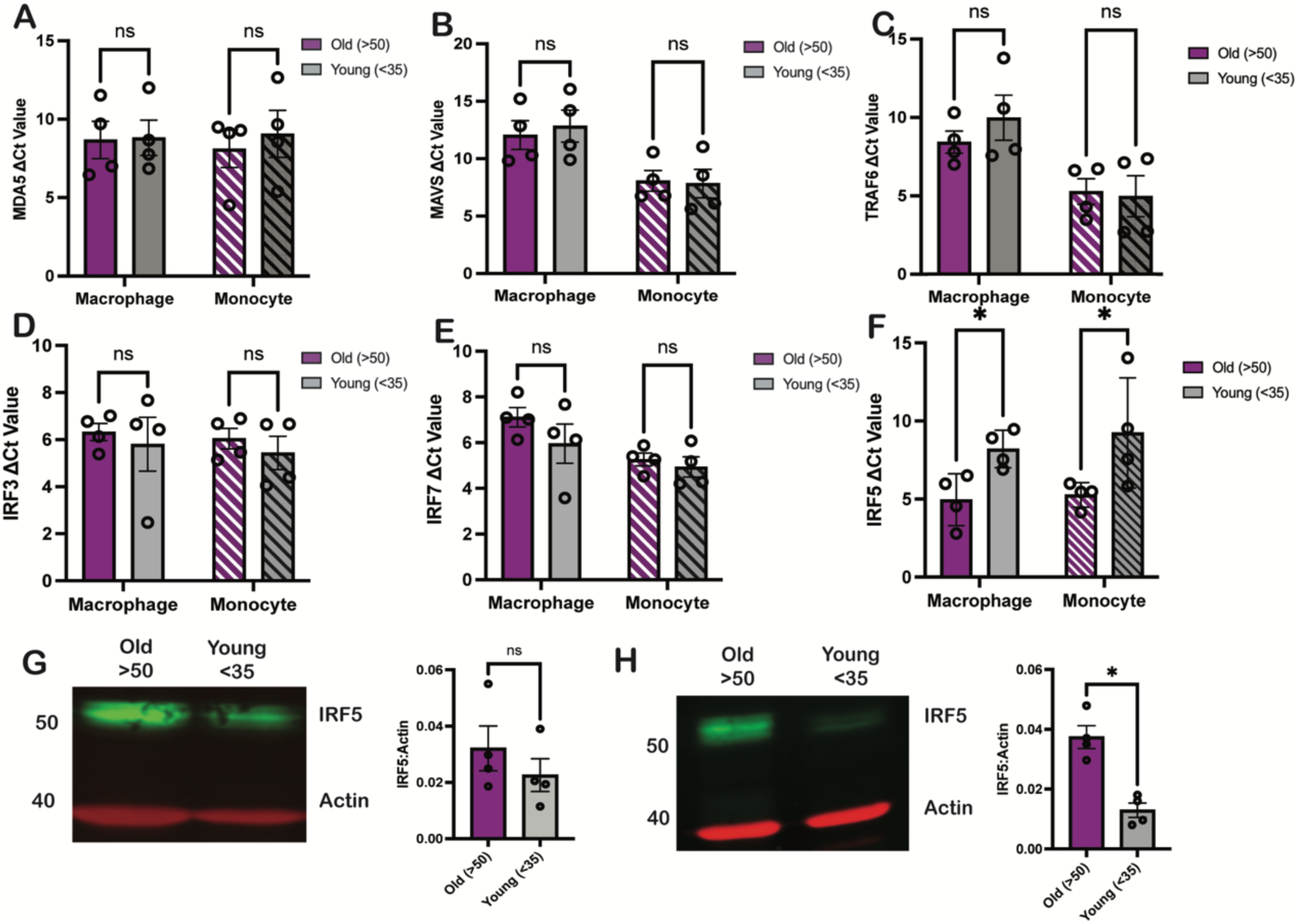
MDMs and monocytes isolated from older individuals exhibit higher levels of IRF5 expression. (A-F) Constitutive mRNA expression of (A) MDA5, (B) MAVS, (C) TRAF6, (D) IRF3, (E) IRF7, and (F) IRF5 in MDMs and monocytes was assessed via RT-qPCR. (G-H) Western blot analysis for constitutive IRF5 expression in MDMs and monocytes. Data is displayed as mean ± SEM with each dot representing a donor. Statistical significance assessed via unpaired I-tests (A-H). *: p < 0.05; ns = not significant.

**TABLE 1.**
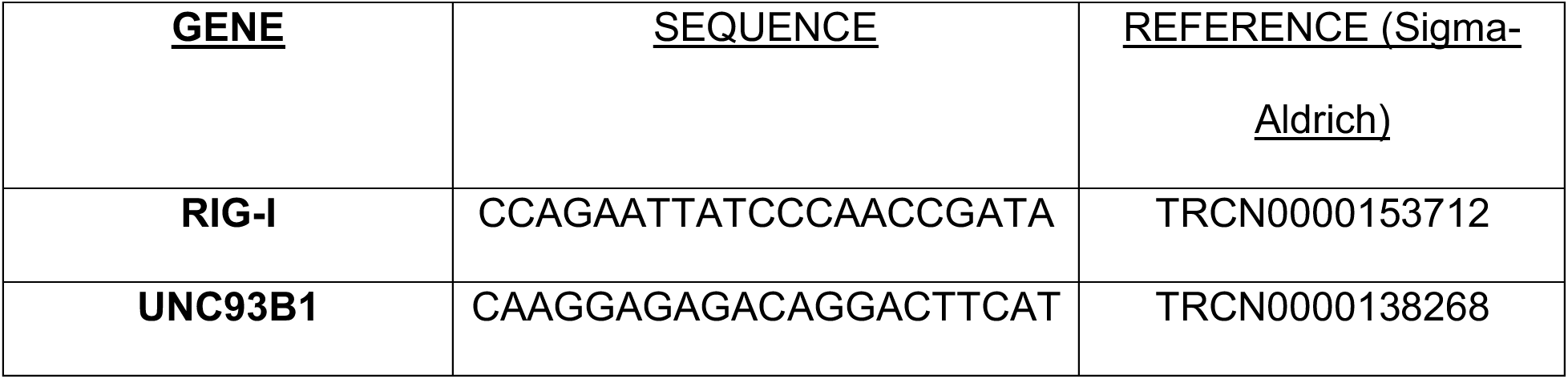
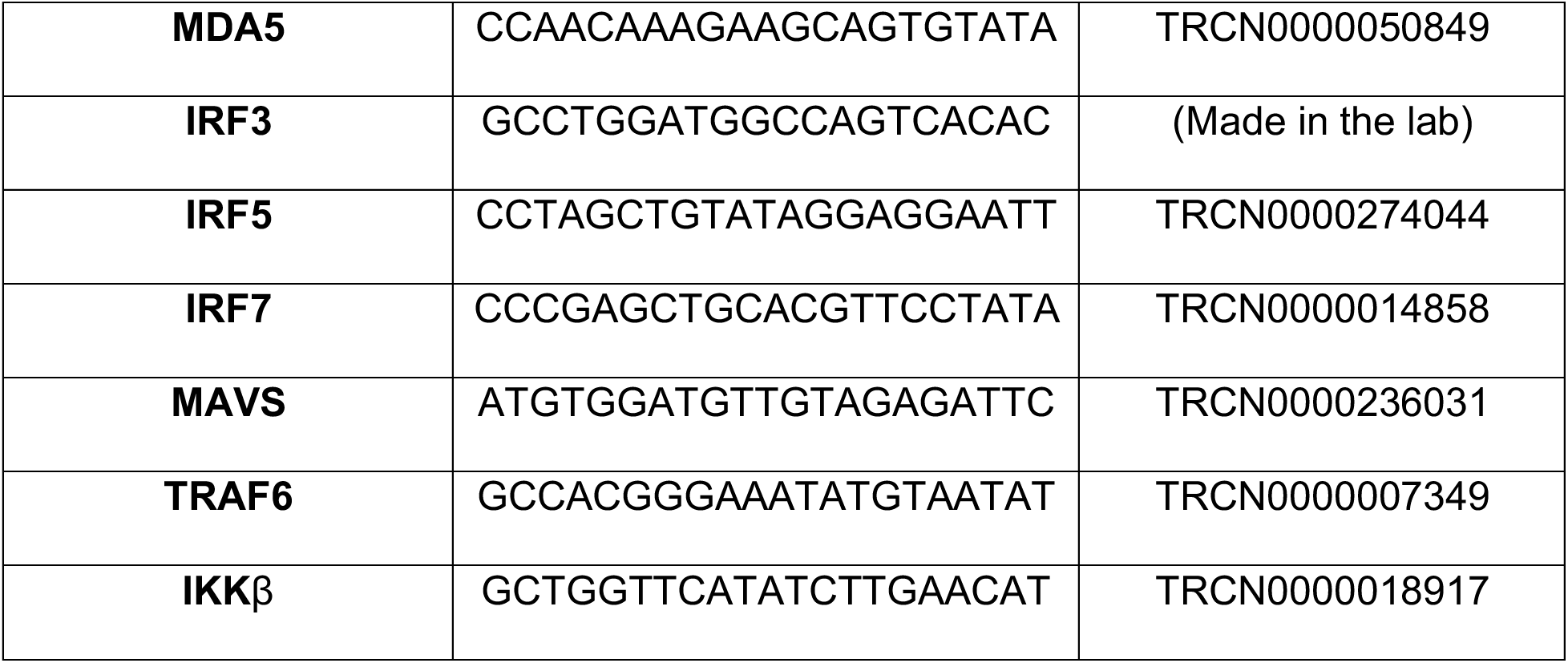

**TABLE 2.**

| **Gene** | **Catalog #** |
| --- | --- |
| RIG-I | L-012511-00-0005 |
| UNC93B1 | L-014650-00-0005 |
| MDA5 | L-013041-00-0005 |
| IRF3 | L-006875-00-0005 |
| IRF5 | L-011706-00-0005 |
| IRF7 | L-011810-00-0005 |
| IKKβ | L-003503-00-0005 |
| TRAF6 | L-004712-00-0005 |

### Innate immune sensing of HIV-1 infection in macrophages is impacted by age

Since IRF5 expression was elevated in macrophages from older donors, we next sought to assess age-related differences in innate immune response to HIV-1 infection. Total RNA was extracted from uninfected and HIV-infected MDMs from 12 donors from the BMC HIV/Aging cohort, and mRNA expression profiles were analyzed via Nanostring using the Myeloid Innate Immunity Panel consisting of 730 target genes. The samples were selected to ensure the same number of older/younger donors (6 each), equal numbers of male/female individuals within each group (3 each per group). We first compared differences in these myeloid innate immune genes at the baseline level in uninfected older and younger MDMs. We observed elevated mRNA levels of several IRF5 target genes including CCL21, IP-10, IL-6, and IL12A/B in older MDMs (Sup Fig. 6A). We also observed slightly elevated expression of IRF5 in MDMs from older donors but did not observe this trend with IRF3 and IRF7 (Sup Fig. 6B). We repeated this analysis using the virus only conditions and found that IRF5, but not IRF3 or IRF7, was significantly upregulated in HIV-1-infected MDMs isolated from older donors (Fig. 8A-B) despite similar levels of infection in vitro (Sup Fig. 6C). We next sought to assess age-related differences in induction of the innate immune responses in older and younger MDMs upon HIV-1 infection. To do this, we compared the fold change in innate immune gene expression in the HIV-1-infected samples to the efavirenz control within each age group. We found that the number of upregulated genes as well as the extent of induction of innate immune gene expression varied based on age. MDMs from older donors demonstrated approximately 97 significantly upregulated genes compared to 42 genes from younger donors (Fig. 8C). These upregulated genes were primarily ISGs including IRF5 and IRF7, as well as several known IRF5 target genes (Fig. 8C) including CCL5, CXCL16 and IL23A, suggesting that MDMs from older individuals display an enhanced IRF5-dependent innate immune response to HIV-1 infection. Additionally, while we observed that IP-10 was upregulated in both older and younger MDMs upon HIV-1 infection, the extent of upregulation was greater in older MDMs (Fig. 8C-D). Surprisingly, we observed no significant induction of IRF5-regulated target gene expression in MDMs from younger donors upon HIV-1 infection (Fig. 8D). Instead, expression of IRF5-regulated genes IL1A and IL23A were significantly down-regulated in response to HIV-1 infection in MDMs from younger donors (Fig. 8D).

**Figure 8:**
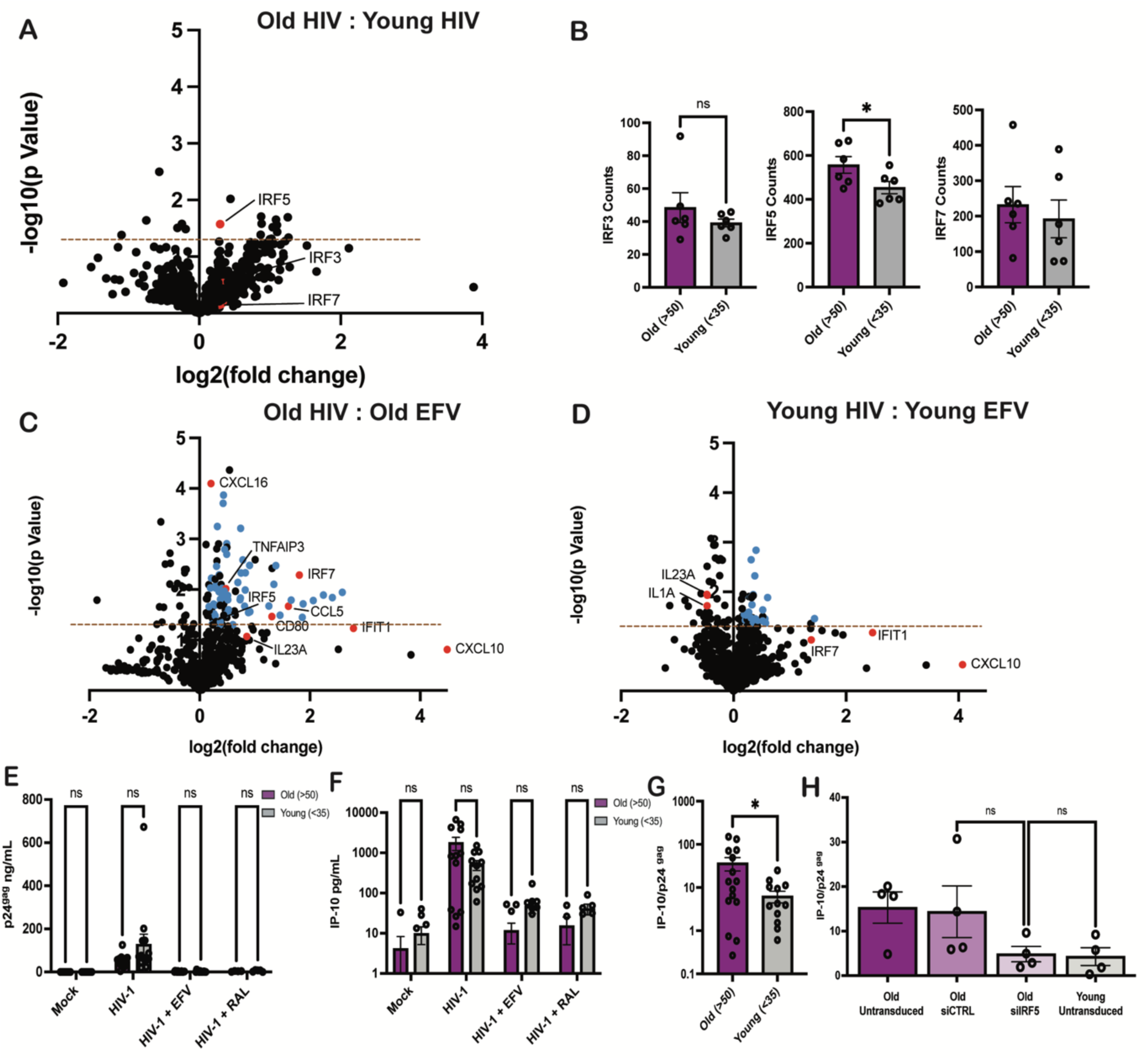
MDMs isolated from older exhibit higher levels of HIV-1 induced immune activation. (A) RNA isolated from Lait.envGFP/G-infected MDMs (MOI 2) was analyzed via Nanostring nCounter using the Myeloid Innate Immunity V2 panel. Baseline expression of the gene panel was calculated using nSolver and plotted as a ratio of Old (HIV} vs. Young (HIV} with IRFs (red} highlighted. The dashed line represents p-value of 0.05. (B) Raw count values for IRF3, IRF5, and IRF7 were plotted to assess differences. (C-D) Fold changes in innate immune gene expression in HIV-infected MDMs ± EFV were plotted. Data from MDMs from (C) older (>50 yo) and (D) younger (<35 yo) donors are shown. Differentially expressed ISGs (blue) and IRFs/lRF targets (red) are highlighted. The dashed line represents a p-value of 0.05. (E-F) Supernatants were analyzed for (E) p24gag production and (F) IP-10 levels via ELISA. (G) IP-10 levels were normalized to those of p24gag for each donor. (H) MDMs were transfected with either control or IRF5 targeting siRNA prior to infection with Lait.envGFP/G. IP-10 and p24gag secretion was measured at 3 dpi. Data is represented as mean ± SEM with each dot representing an individual donor (B, E-G) or with each dot representing a target gene (A, C-D). Significance was assessed via unpaired two-tailed I-test (A-B, E-G}, paired two-tailed I-test (C-D}, or 1-way ANOVA with Tukey’s multiple comparisons test (H). *: p<0.05, ns = not significant.

Interestingly, while there was no significant difference in extent of HIV-1 infection in MDMs derived from younger or older MDMs (Fig. 8E and Sup Fig. 5A), IP-10 production in HIV-infected older MDMs was higher than that observed with HIV-infected MDMs from younger donors (Fig. 8F and Sup Fig. 5B). The difference was evident even when IP-10 secretion levels were normalized to HIV-1 infection levels (p24 production in the culture supernatants) across donors to account for donor-to-donor variation in infection establishment (Fig. 8G). In contrast, no significant age-associated differences in IP-10 production was observed upon stimulation with 3p-hp RNA (RIG-I agonist) or LPS (TLR4 agonist) in MDMs (Sup. Fig. 5C-F), suggesting an HIV-specific enhanced innate immune response in macrophages from older donors. Finally, knock-down of IRF5 expression in older MDMs suppressed HIV-infection induced IP-10 secretion to levels observed in younger MDMs (Fig. 8H). Taken together, these results indicate that the inflammatory response induced upon MDA5 sensing of HIV-1 icRNA expression in MDMs is impacted by age and cell-intrinsic IRF5 expression. Importantly, constitutively elevated IRF5 expression in macrophages from older donors might contribute to the heightened pro-inflammatory state and accelerated course of clinical disease in older PWH.

## Discussion

In this study, we highlight an important, non-redundant role of MDA5 in sensing of HIV-1 icRNA in macrophages and induction of type I IFN responses. These findings were corroborated in a recent study that showed MDA5-mediated sensing of HIV-1 icRNA in dendritic cells (57, 58). Though the specific motif within HIV-1 icRNA that is recognized by MDA5 is currently unclear, sensing of non-self RNA by RLR family members is dependent on unique invariant features. For instance, discrimination between cellular and viral dsRNA by RIG-I is based on 5’ tri-phosphate motifs and blunt-ended RNA duplex structure of <100bp in length, though size requirement can be variable (70). Though previous studies suggest that transfected HIV-1 RNA can be sensed by RIG-I, it is unclear whether this mechanism is conserved in infection models (71). Importantly, in our studies, knock-down of RIG-I expression in macrophages had no impact on HIV-1 icRNA-induced type I IFN responses (Fig. 2). In contrast, MDA5 does not require terminal 5’-tri-phosphates for ligand recognition, but rather preferentially recognizes long dsRNAs (>500bp), and utilizes length specificity to distinguish self from non-self dsRNAs (72, 73). Ligand binding nucleates MDA5 filaments on dsRNA, and subsequent recruitment and induction of MAVS filament formation in a caspase activation and recruitment domain (CARD)-dependent manner(74).

Our results demonstrate specific immunoprecipitation of HIV-1 icRNA but not msRNA by MDA5 in virus-infected cells. While, MDA5 is known to bind to long dsRNA genomes or replication intermediates of single strand viruses (75), it is plausible that MDA5 recognizes duplex regions of stem-loops (76, 77) or other uncharacterized RNA features uniquely existing in HIV icRNA. Further studies are warranted to elucidate the molecular mechanisms underlying MDA5 sensing of HIV icRNA. Interestingly, MDA5 has also been implicated in sensing of endogenous retroelement 3’ UTR-containing dsRNAs generated either via bidirectional transcriptional mechanisms or upon cytoplasmic spillover of intron-retained endogenous retroelement (ERE) RNAs (78). While expression of ERE RNAs is enhanced due to transcriptional de-repression in senescent or aging cells (79) or exacerbated due to HIV-1 infection (80, 81), contributions of ERE dsRNAs to innate immune activation in HIV-infected cells need further investigations. Additionally, while other studies have implicated TLR7 in sensing of HIV-1 gRNA in plasmacytoid dendritic cells (82), abrogation of endosomal TLR sensing in macrophages upon downregulation of UNC93B1 expression did not impact HIV-1 icRNA-induced IP-10 and IFNβ induction, suggesting that the mechanism of HIV-1 RNA sensing varies in a cell-type dependent manner, and that MDA5 is required for sensing of intron-containing HIV-1 transcripts in macrophages.

This study also highlights an important role of IRF5 in the innate immune signaling pathway downstream of MDA5/MAVS-dependent HIV-1 icRNA sensing. While previous studies have implicated IRF3 and IRF7 in inducing expression of IFNβ and ISGs in HIV-1-infected cells (83–86), here we show that IRF5 also has an important role in inducing IFNβ and pro-inflammatory cytokine IP-10 expression in HIV-1-infected macrophages, suggesting functional redundancies in IRF requirements. This is consistent with previous findings that IRF5 does not bind to virus response elements (VREs) in the promoter region of IFNα, which highlights that IRF5 activation leads to the transcription of specific type I IFNs and ISGs (87). This IRF5 DNA binding specificity may also account for conflicting findings that IRF7, but not IRF5, is crucial for ISG15 induction in HIV-1-infected macrophages and DCs (57, 58). We also find that the type of viral infection has an impact on IRF5 involvement in mediating IP-10 production. For instance, IRF5 knockdown attenuated IP-10 production in HSV-1 but not in Sendai virus infected cells, highlighting that IRF5 activation might uniquely be dependent on the infecting virus and their replication intermediates.

Previous studies have shown that MAVS activation downstream of RLR sensing pathways leads to recruitment of various E3 ubiquitin ligases, TRAF2, TRAF5, and TRAF6, and cytosolic serine kinases, IKK and TBK1 (88, 89) for IFNβ induction. As opposed to the redundant roles of TRAF2/3 and TRAF6 for NF-κB and IRF3 activation and IFNβ induction downstream of Sendai virus infection (88), our results suggest a non-redundant role of TRAF6 in IRF5 activation downstream of HIV-1-icRNA sensing in macrophages. These findings align with the purported requirement for TRAF6-mediated K63-linked ubiquitination of IRF5 as an essential protein modification for IRF5 activation (63). Interestingly, knock-down of IKKβ abrogated IP-10 and IFNβ induction in HIV-1-infected macrophages, highlighting the essential role for additional post-translational modifications of IRF5 such as phosphorylation (64) (66, 90) downstream of HIV-1 icRNA sensing by MDA5 and MAVS. Taken together, we propose that HIV-1 icRNA sensing by MDA5 and MAVS activation leads to the recruitment and activation of TRAF6 and IKK∆, resulting in ubiquitination and phosphorylation and subsequent nuclear translocation of IRF5, contributing to IP-10 and IFNβ expression. IFNβ production might further induce IRF5 expression thus exacerbating production of type I IFNs and proinflammatory cytokines.

As the population of individuals living with HIV increases in age, it is crucial to understand the mechanisms that contribute to higher levels of immune activation in older PWH (91–93). While duration of ART, sex, lifestyle and comorbidities are all contributing factors, results described in this study propose a cell-intrinsic role of IRF5 in inducing innate immune activation in HIV-1-infected macrophages from older donors. We found that MDMs and monocytes from older donors in two distinct cohorts had higher baseline levels of IRF5 expression at the mRNA and protein level, and consequently expressed elevated mRNA expression levels of several IRF5 target genes including CCL21, IP-10, IL12A, IL12B, and IL-6. Upon HIV-1 infection of MDMs, we also observed a higher level of ISG induction in MDMs from older donors, including IRF5, as well as several known IRF5-responsive genes such as CXCL16, CCL5, and CD80. IP-10 has been shown to be upregulated in the plasma of older individuals and is negatively correlated with working memory (94). Serum IL-6 levels also increase with age and have been associated with frailty and mortality in chronic disease settings (95–97) as well as age-related neurodegenerative diseases such as Alzheimer’s (98–100). Additionally, significantly higher serum levels of CXCL16 were correlated with severe COVID19 outcome, which is more common among individuals over 65 years old (101, 102). Similar to previous studies which reported elevated IRF5 expression in monocytes from older donors compared to younger donors upon RIG-I activation (103), MDA5/MAVS-mediated IRF5 nuclear translocation in HIV-1-infected macrophages and type I IFN-induced IRF5 expression might contribute to IRF5 hyperactivation. Taken together, these results indicate that enhanced levels of constitutive IRF5 expression in older monocytes ad macrophages and HIV-infection-induced IRF5 activation may contribute to enhanced inflammatory responses in PWH and persistent expression of “inflammaging”-related genes.

IRF5 has been considered a promising therapeutic target for a variety of inflammatory diseases such as systemic lupus erythematosus and rheumatoid arthritis (104, 105) IRF5 function is cell-type specific and limited primarily to induce pro-inflammatory cytokines in response to infection (106, 107). Unlike other transcription factors such as NF-κB, the downstream targets of IRF5 are more specific (108, 109), and incorporation of IRF5-targeting therapeutics in combination with ART could potentially reduce chronic inflammation and thus prevent the development of co-morbidities associated with accelerated inflammaging in older PWH.

## Methods

### Study Population

PBMCs were derived from the HIV and Aging Cohort established at Boston Medical Center (BMC HIV Aging cohort) (26). Cohort recruited people without HIV in two age-stratified groups, younger (18-35 years old) and older (aged ≥50 years old) group. Donors with active hepatitis B or C, or recent immunomodulatory therapy (oral or injected corticosteroids, plaquenil, azathioprine, methotrexate, biologic therapies, systemic or local interferon, chemotherapy or HIV vaccine) were excluded (26). PBMCs were also isolated from leukopaks (NY Biologics, Long Island, NY) from anonymous donors stratified by age into younger (18-35 years) or older (≥50 years).

### Plasmids

Single-cycle HIV-1 encoding GFP in place of *nef* (LaiΔenv/GFP) and an HIV-1 Rev mutant (M10) deficient in CRM1 binding (LaiΔenv/GFP-M10) have been previously described (44, 110). Lentiviral vectors (pLKO.1) expressing shRNAs against IRF3, IRF5, IRF7, RIG-I, UNC93B1, MDA5, TRAF6, and IKK∆ were purchased from Sigma-Aldrich (or constructed by ligating annealed double-stranded oligonucleotides into pLKO.1 using AgeI and EcoRI sites). HIV-1 packaging plasmid, psPAX2, and VSV-G expression plasmid, H-CMV-G, have been described previously (111).

### Cells

HEK293T cells (ATCC) and TZM-bl (NIH AIDS Reagent Program) were cultured in DMEM (Gibco) supplemented with 10% fetal bovine serum (FBS) (Gibco) and 1% penicillin/streptomycin (pen/strep) (Gibco). THP-1 cells (NIH AIDS Reagent Program) were cultured in RPMI (Gibco, 11875-119) supplemented with 10% FBS and 1% pen/strep (R10). THP-1 cells were differentiated to macrophages with 100nM PMA (Sigma-Aldrich) for 48 hours as described previously (56). THP-1 cells were transduced with lentivectors (100 ng p24^gag^ per 0.5x10^6^ cells) expressing shRNAs (shControl, shMDA5, shMAVS, shTRAF6, shIKKβ, shIRF3, shIRF5, and shIRF7). Sequences of shRNA targets are listed in Table 1. Transduced cells were maintained and selected in the presence of 2 µg/mL puromycin (Invivogen). Validating knockdown cell lines at the functional level were carried out by incubating PMA/THP-1 cells with a variety of RLR or TLR ligands for 24-48 hours and measuring IP-10 in the culture supernatants via ELISA (described below). Ligands used include 3p-hpRNA (2.5 ng/mL, Invivogen tlrl-hprna), LPS (TLR4 ligand, 100 ng/mL, Invivogen tlrl-eblps), or high molecular weight (HMW) poly I:C (TLR3 ligand, 10 ug/mL, Invivogen, tlrl-pic). Human monocyte derived macrophages (MDMs) were derived from CD14+ monocytes isolated from PBMCs by anti-CD14 antibody conjugated magnetic beads (Miltenyi Biotech) and cultured in RPMI supplemented with 10% Human AB Serum (Sigma-Aldrich) and 20 ng/ml M-CSF (Peprotech,) for 6 days as previously described (44, 56).

### Viruses

VSV-G-pseudotyped single-round HIV-1 were generated by transient transfection of HEK293T cells as previously described (44). Virus titer was measured in TZM-bl cells (112). Lentivectors expressing shRNA were generated by co-transfecting HEK293T cells with pLKO.1, psPAX2 and H-CMV-G using calcium phosphate (111). Virus containing supernatants were harvested 2 days post-transfection, passed through 0.45 µm filters and concentrated on a 20% sucrose cushion (24,000 rpm at 4 °C for 2 h with a SW28 rotor (Beckman Coulter)). Virus pellets were resuspended in 1xPBS, aliquoted, and stored at -80 °C until use. Lentiviral p24^gag^ content was measured via p24^gag^ ELISA as previously described (111). HSV-1 and Sendai viruses were generously provided by Dr. Mohsan Saeed (Boston University CAMed).

### Infections

THP-1/PMA macrophages were infected with single-cycle HIV-1 reporter viruses (MOI 2) in presence of polybrene (10 μg/mL, Millipore Sigma, TR-1003-G) via spinoculation (2,300 rpm, 1h at RT). In some experiments, THP-1/PMA macrophages were pre-treated (for 20 minutes) with efavirenz (1 µM, NIH AIDS Reagent Program), raltegravir (30 µM, Selleck Chemical #50-615-1), spironolactone (100 nM, Selleck Chemical, # S4054), and KPT 330 (1 µM, Selleck Chemical # 50-136-5156). MDMs were pre-treated with 2.5 mM dNs (Sigma-Aldrich)(113–115) for 2 hours and then infected with LaiΔenvGFP/G or LaiΔenvGFP-M10/G (MOI 1) as previously described (44) in the presence or absence of HIV-1 inhibitors.

### siRNA transfection of primary MDMs

MDMs (1.5x10^6^ cells/well in a 6-well plate) were transfected with SMARTPool siRNA (25-50 nM, Horizon) via Trans-IT X2 (Mirius Bio, MIR6004) in Opti-MEM (Gibco). SMARTPool siRNA catalog numbers are listed in Table 2. Transfected cells were detached with enzyme-free cell dissociation buffer (Millipore, # S014B) at 24 h post transfection and reseeded for infections as described above.

### RNA Isolation and RT-qPCR

Total RNAs were extracted using the RNeasy kit (Qiagen). cDNA was generated using Superscript III 1^st^ Strand cDNA Synthesis kit (Invitrogen), and qPCR was done using Maxima SYBR Green (Thermo Scientific). C_T_ value of target mRNA was normalized to that of GAPDH mRNA (ΔC_T_), then ΔC_T_ of the target mRNA was further normalized to that of a control sample by the 2-^ΔΔC^_T_ method as described (116, 117). Primer sequences for GAPDH, IP-10, IFNα, and IFNβ have been described previously (44). Primer sequences to assess expression of RIG-I. MDA5, UNC93B1, IRF3, IRF5, IRF7, MAVS, TRAF6 and IKKβ are listed in Table 3.

**TABLE 3.**
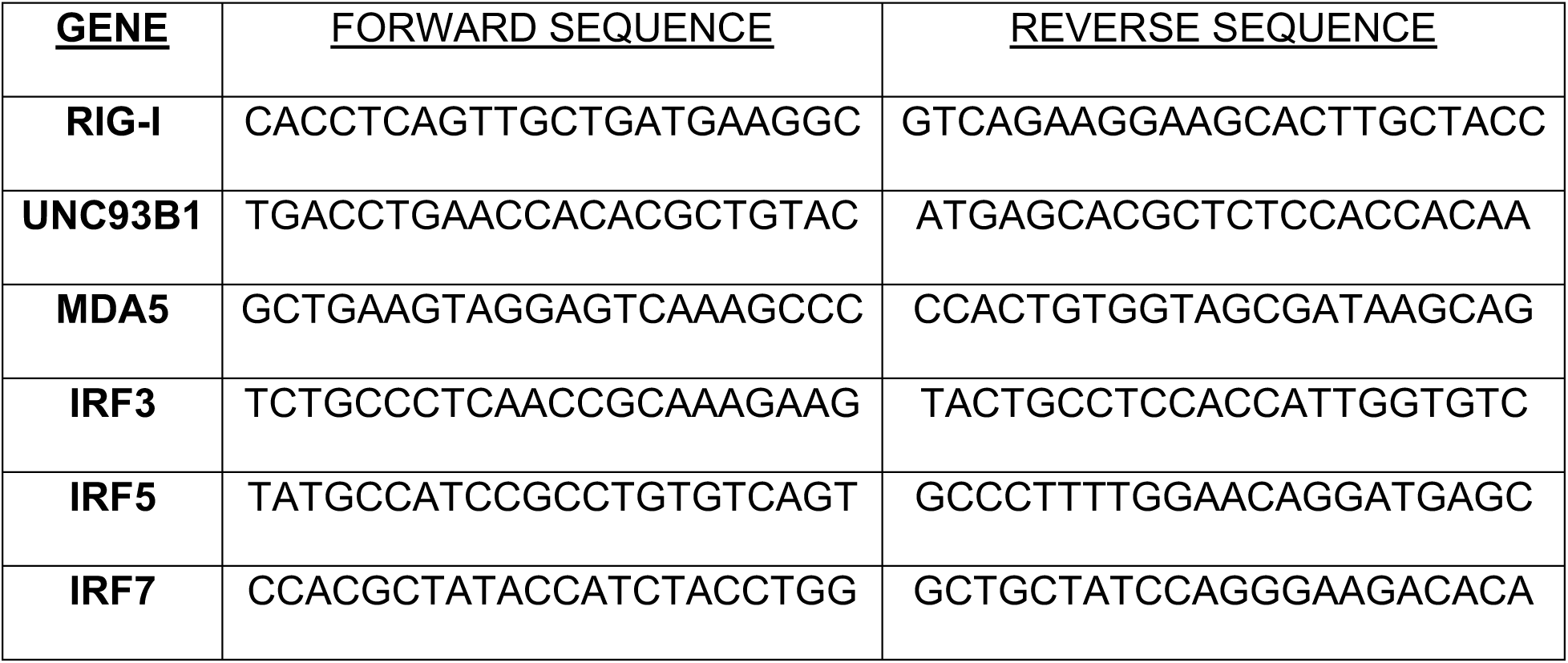
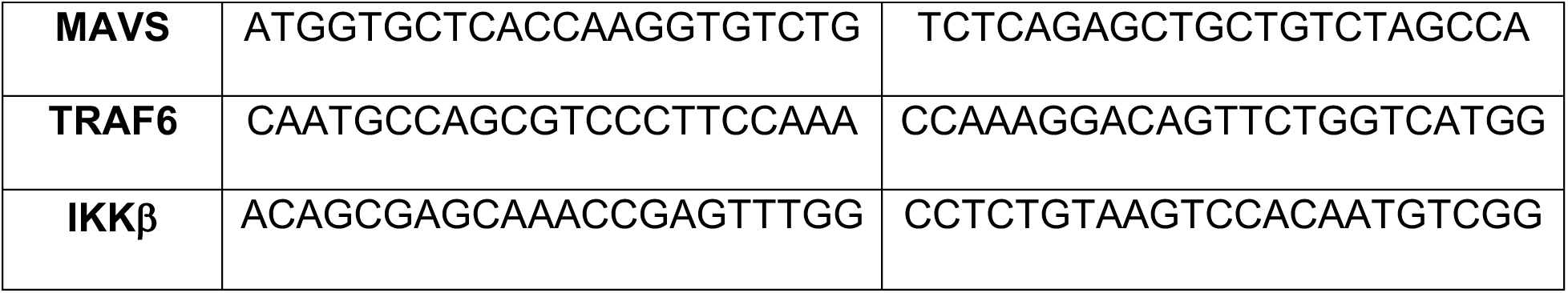

### Flow Cytometry

THP-1/PMA macrophages were detached using Enzyme Free Dissociation Buffer (Millipore, S014B), while MDMs were detached using CellStripper (Corning, MT-25-056CI), washed once with 1x PBS, and fixed in 4% PFA for at least 30 minutes. Cells were analyzed on LSRII (BD) flow cytometer to assess infection efficiency via GFP positivity. Data was analyzed using FlowJo software (FlowJo).

### Western Blot analysis

In order to assess levels of IRF5 expression or validate knockdown of host factors, cell lysate (30 μg) were analyzed by western blotting using the following antibodies: rabbit anti-MAVS (Invitrogen, PA5-17256, 1:1000), rabbit anti-IRF3 (Cell Signaling, 4302S, 1:1000), rabbit anti-IRF5 (Cell Signaling, 76983S, 1:1000), rabbit anti-IRF7 (Cell Signaling, 4920S, 1:1000), rabbit anti-TRAF6 (Cell Signaling, #8028, 1:1000), rabbit anti-IKKβ (Cell Signaling, 2684S, 1:1000), mouse anti-actin antibody (Thermo Fisher, #AM4302, 1:5000), goat anti-mouse IgG secondary antibody Dylight 680 (Thermo Fisher, SA5-35518, 1:10,000), and goat anti-rabbit IgG secondary antibody Dylight 800 (Thermo Fisher, SA5-35571, 1:10,000). Membranes were scanned using an Odyssey scanner (Li-Cor).

### Immunofluorescence and Microscopy

THP-1/PMA macrophages were cultured and infected on coverslips (Fisher Scientific, 12-541-001) in 12-well plates. Three days post-infection, cells were fixed using 4% PFA (BM-155, Boston Bioproducts) for 30 min at 4°C, washed once with PBS and permeabilized using 0.1 % Triton X-100 (NC1636886, Cayman Chemical) in PBS for 5 minutes at room temperature. Cells were stained with anti-IRF5 antibody (Cell Signaling, 76983S) and Alexa594-conjugated goat anti-mouse antibody (Invitrogen), and counterstained with DAPI (Sigma-Aldrich, D1306). Cells were mounted on slides using Fluoromount mounting medium (Southern Biotech, OB100-01). Images were acquired using an EVOS M5000 Microscope (Invitrogen, A40486) or a Leica SP5 Confocal Microscope and analyzed using ImageJ (NIH) or CellProfiler (118, 119).

### RNA Co-Immunoprecipitation Assay

HEK293T cells were infected with Lai∆envGFP/G at MOI 1, and transfected 18 h post infection with either 3 μg of MDA5-Flag plasmid or 0.1 μg of RIG-I-Flag plasmid (+ filler plasmid). The next day, cells were washed twice with PBS and UV crosslinked at 40 mJ/cm^2^. Cells were lysed with 400 μL of Fractionation Buffer (Life Technologies, 4403461) supplemented with protease inhibitor (Roche, 11836170001), RNAseOUT (100 U/ml, Invitrogen, 10777019), and DTT (5 µL/ml, Invitrogen, 18080051). 40 µL of lysate was saved as ‘input’ and the remaining cytoplasmic lysate was incubated with antibody coupled beads as described previously (120). Beads (Invitrogen, 61-011-LS) were coated with 2µg of anti-Flag antibody (Sigma, F3165) or IgG control (Novus Biologicals, NBP1-97019). Following overnight antibody coupling, beads were washed using high salt buffers as described previously (121). RNA extraction, cDNA synthesis, and qPCR were carried out as previously described (120).

### ELISA

IP-10 in THP-1/PMA and MDM supernatants was measured with ELISA according to manufacturer’s protocol (BD Biosciences, 550926).

### Nanostring Analysis

Total RNA was isolated from infected MDMs using RNeasy kit (Qiagen, #74106) and quantified via Nanodrop. Samples were analyzed using the Human Myeloid Innate Immunity V2 panel (Nanostring, 115000171) on the NCounter system. Nanostring counts were analyzed using nSolver; raw counts of all targets normalized to geometric mean of positive controls, housekeeping controls included in the Nanostring Myeloid Innate Immunity panel. Counts between donors were compared using two-tailed t-test and data was plotted as log_2_(fold change) vs -log(p-value) with the line representing p-value of 0.05.

### Statistics

Statistical analysis was performed using GraphPad Prism 10. P-values were calculated via 1-way ANOVA with Tukey’s post-test or Dunnett’s post-test for multiple comparisons analysis, or unpaired t-test.

## Acknowledgements

We thank the HIV/Aging Cohort (BMC) study participants as well as the Boston Medical Center Infectious Disease Clinical Research Unit. We are grateful to Dr. Mohsan Saeed (BUMC) for gracious gifts of HSV and Sendai virus stocks. We also thank the BUMC Flow Cytometry Core and Cellular Imaging Core for technical assistance. This work was supported by R01AG060890 (S.G. and M.S.), R01DA051889 (S.G.), R01DA055488 (S.G. and A.H.) and P30AI042853 (S.G. H.A., A.H., and M.S.).

## Author contributions

S.R., H.A. and S.G. designed the experiments. S.R. A.Q. and J.B. performed the experiments and analyzed the data. A.O., Y.C., Y.L., A.A., and M.S. assisted in obtaining and isolating PBMCs from participants of the HIV/Aging Cohort (BMC). S.R., H.A., and S.G. wrote the manuscript.

---

**Supplementary Table 1.**

| Group | Old (>50) | Young (<35) |
| --- | --- | --- |
| Age (Mean ± SD) | 64 ± 7.9 | 29.5 ± 4.2 |
| Sex (% Male) | 50 | 50 |
| BMI (Mean ± SD) | 28.5 ± 2.9 | 30.5 ± 7.9 |
| Gastrointestinal Disorder (%) | 66.7 | 16.7 |

**Supplementary Table 2.**
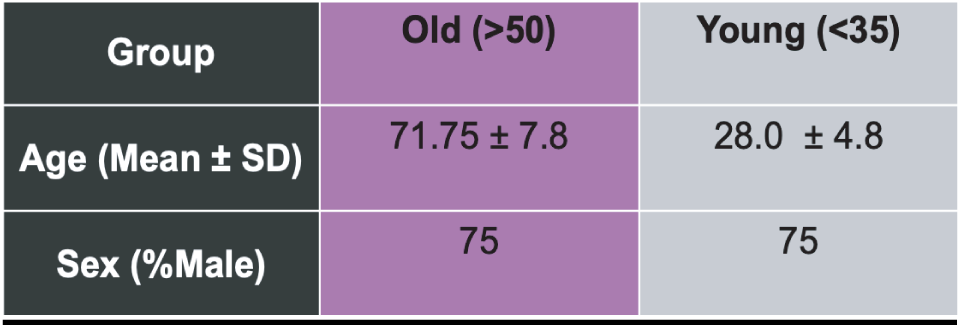

| Group | Old (>50) | Young (<35) |
| --- | --- | --- |
| Age (Mean ± SD) | 71.75 ± 7.8 | 28.0 ± 4.8 |
| Sex (%Male) | 75 | 75 |

Table 1 summarizes characteristics of participants in the HIV/Aging Cohort (BMC).Table 2 summarizes donor characteristics of leukopaks obtained from NY Biologics.

**Supplementary Figure 1.**
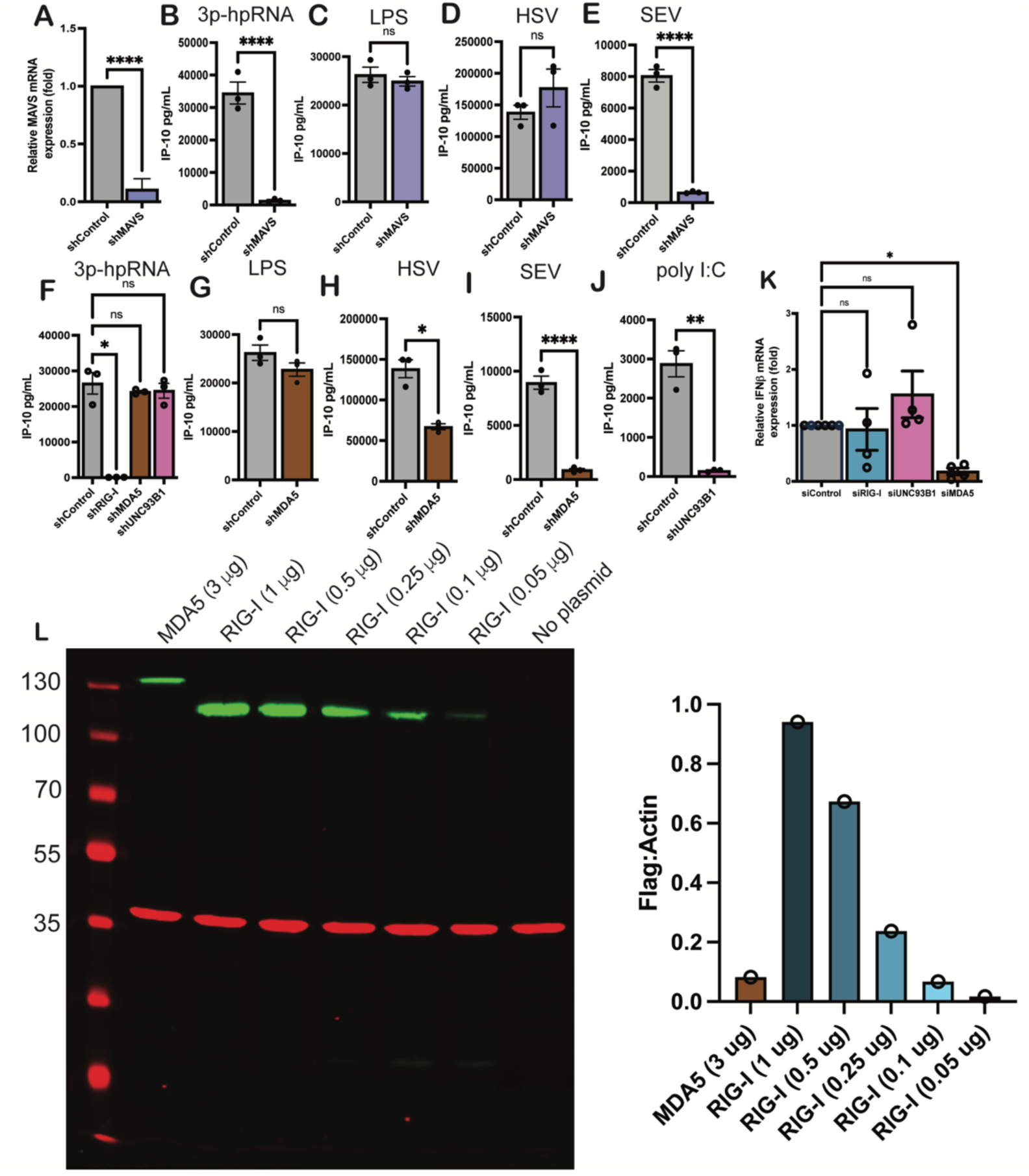
(A) Knockdown of MAVS expression in THP-1/PMA macrophages was validated using RT-qPCR. (B-E) THP-1/PMA cells expressing MAVS shRNA were treated with 3p-hpRNA (2.5 ng/ml), LPS (100 ng/ml), infected with HSV (MOI 0.1) or Sendai virus (SEV) (MOI 1), andsupernatants were harvested 24 h post-stimulation or 48 h post-infection for analysis of IP-10 production via ELISA. (F-J) THP-1/PMA cells with knock-down of RIG-I, MDA5 or UNC93B1 expression were treated with 3p-hpRNA (2.5 ng/ml) (F), LPS (100 ng/ml) (G), HMW polyl:C (10 mg/ml) (J) or infected with HSV (MOI 0.1) (H) or Sendai virus (MOI 1) (I). Supernatants were harvested 24 hours post-stimulation or 48 h post infection for analysis of IP-10 production via ELISA. (K) siRNA-transfected MDMs were infected with Lai6env GFP/G (MOI 1) in the presence of dNs and harvested for analysis of IFN mRNA expression via RT-qPCR at 2 dpi. Data is displayed as mean± SEM with each dot representing an experiment (A-J) or individual donor (K). Statistical significance assessed via unpaired t-test (A-E, F-J,L-N,S-U) or 1-way ANOVA (F, K,O-R,V) with Dunnett’s multiple comparisons analysis.*: p < 0.05; **: p<0.01, ***: p < 0.001 ****: p < 0.0001, ns = not significant. (L) HEK293T cells were transfected with varying concentrations of RIG-I-Flag, or a single amount of MDA5-Flag expression plasmids. Expression of MDA5/RIG-I-Flag was assessed via Western blot analysis. Flag intensity was normalized to that of actin and plotted as a graph.

**Supplementary Figure 2.**
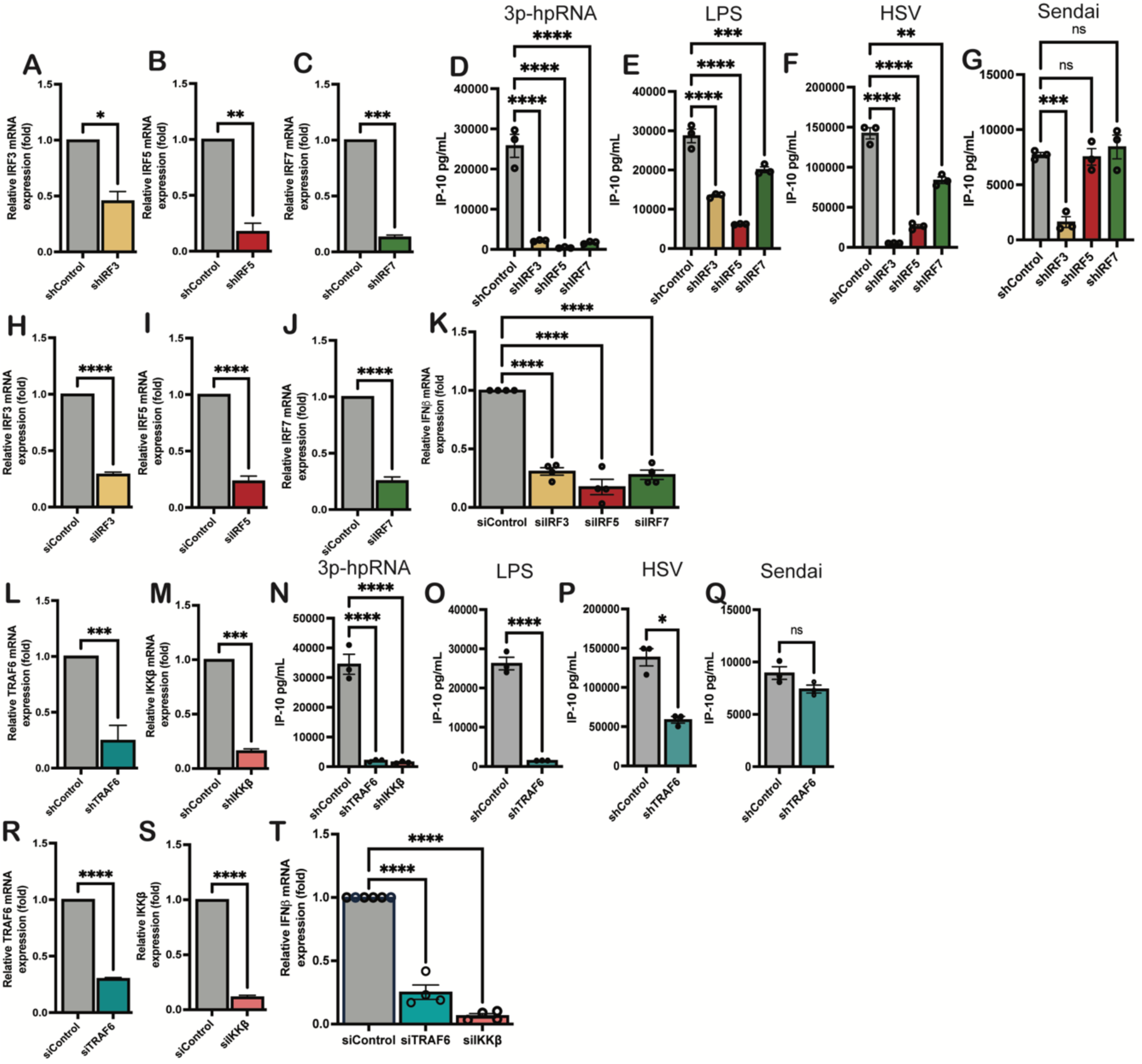
Knockdown of (A) IRF3, (B) IRF5, or (C) IRF7 expression in THP-1/PMA macrophages was validated by RT-qPCR for (n=3). Functional knockdown of IRF3, IRF5 and IRF7 expression in THP1/PMA macrophages was validated using (D) 3p-hpRNA (2.5 ng/ml), (E) LPS (10Ong/ml), (F) HSV (MOI 0.1) infection or (G) Sendai virus (MOI 1) infection. MDMs were transfected with siRNA targeting (H) IRF3, (I) IRF5, or (J) IRF7 for 48 hours and knockdown validated using RT-qPCR (n=4). siRNA transfected MDMs were infected with LaiL’.envGF­ P/G (MOI **1)** in the presence of dNs and harvested for analysis of IFNf3mRNA expression (K) via RT-qPCR at 2 dpi. Knockdown of TRAF6 **(L)** and IKKf3 (M) expression in THP-1/PMA macrophages was validated using RT-qPCR, and functionally validated by treatment with 3p-hpRNA (2.5 ng/ml) (N), LPS (100 ng/ml) (0), infection with HSV (MOI 0.1) (P) or Sendai virus (MOI 1) (Q). Cell supernatants were harvested 24 h post-stimulation or 48 h post infection for analysis of IP-10 production via ELISA. (R-S) MDMs were transfected with SmartPool siRNA against (R) TRAF6 or (S) IKKf3and mRNA expression was measured by RT-qPCR (n=4). (T) MDMs transfected with SmartPool siRNA targeting TRAF6 or IKKf3 and infected with LaiMnvGFP/G (MOI 1) in the presence of dNs and harvested for analysis of IFNf3 mRNA expression at 2 dpi. Data is represented as mean± SEM with each dot representing an individual experiment (A-G, L-Q) or donor (K,T). Statistical significance was assessed using unpaired I-test (A-C, H-J, L-M, O-S) or 1-way ANOVA with Dunnett’s multiple comparisons analysis (D-H, K, N, T). *: p < 0.05; **: p<0.01, ***: p < 0.001 ****: p < 0.0001, ns = not significant.

**Supplementary Figure 3.**
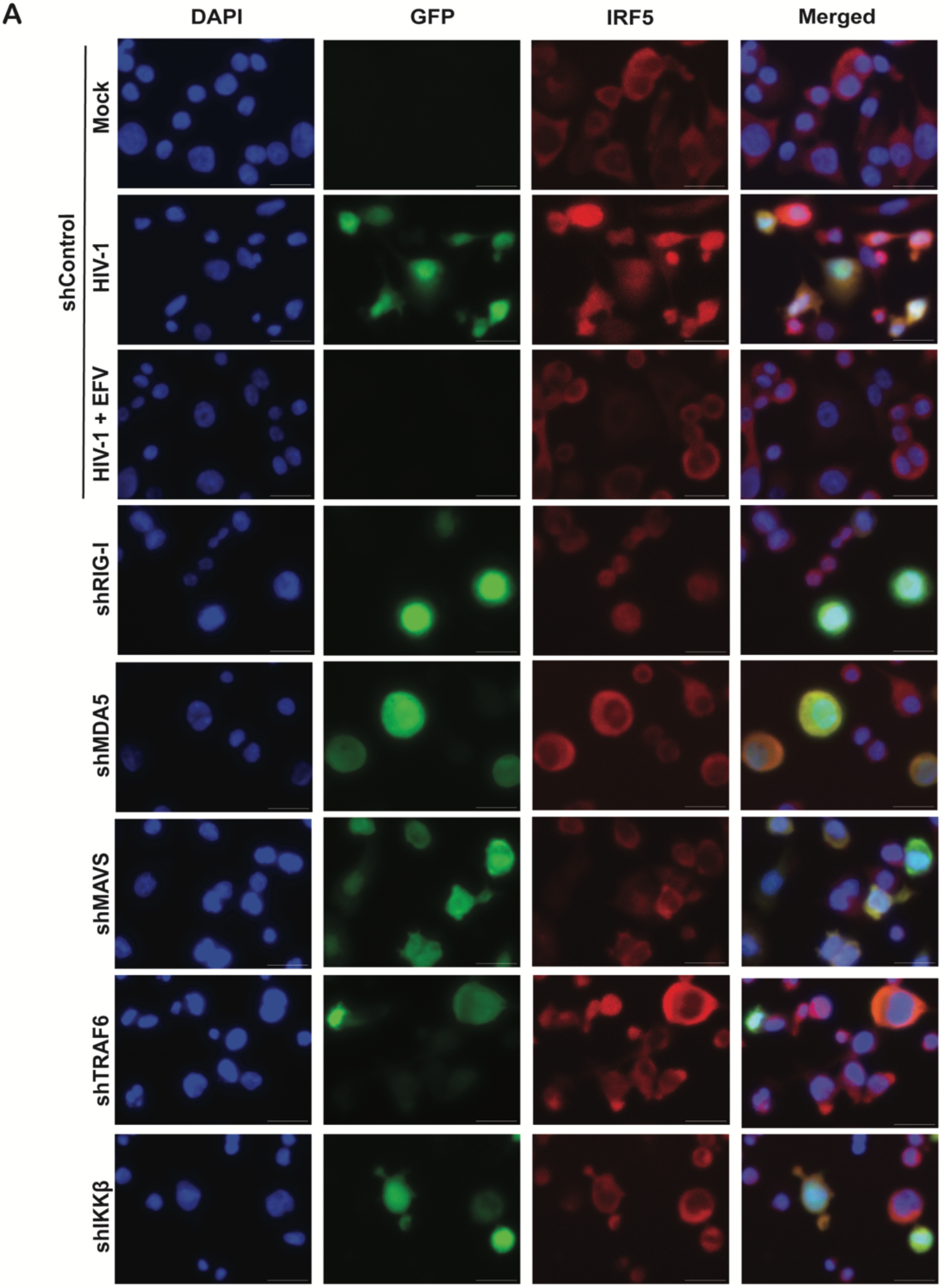
THP1/PMA macrophages transduced with lentivectors expressing Ctrl, RIG-I, MDA5, MAVS, TRAF6 or IKK shRNAs cells were infected with LaiMnvGFP/G (MOI 2) ± EFV (1µM) on coverslips. Cells were fixed at 3 dpi and stained to visualize intracellular IRF5 localization and DAPI via immunofluorescence imaging. Scale bar= 100 µm.

**Supplementary Figure 4.**
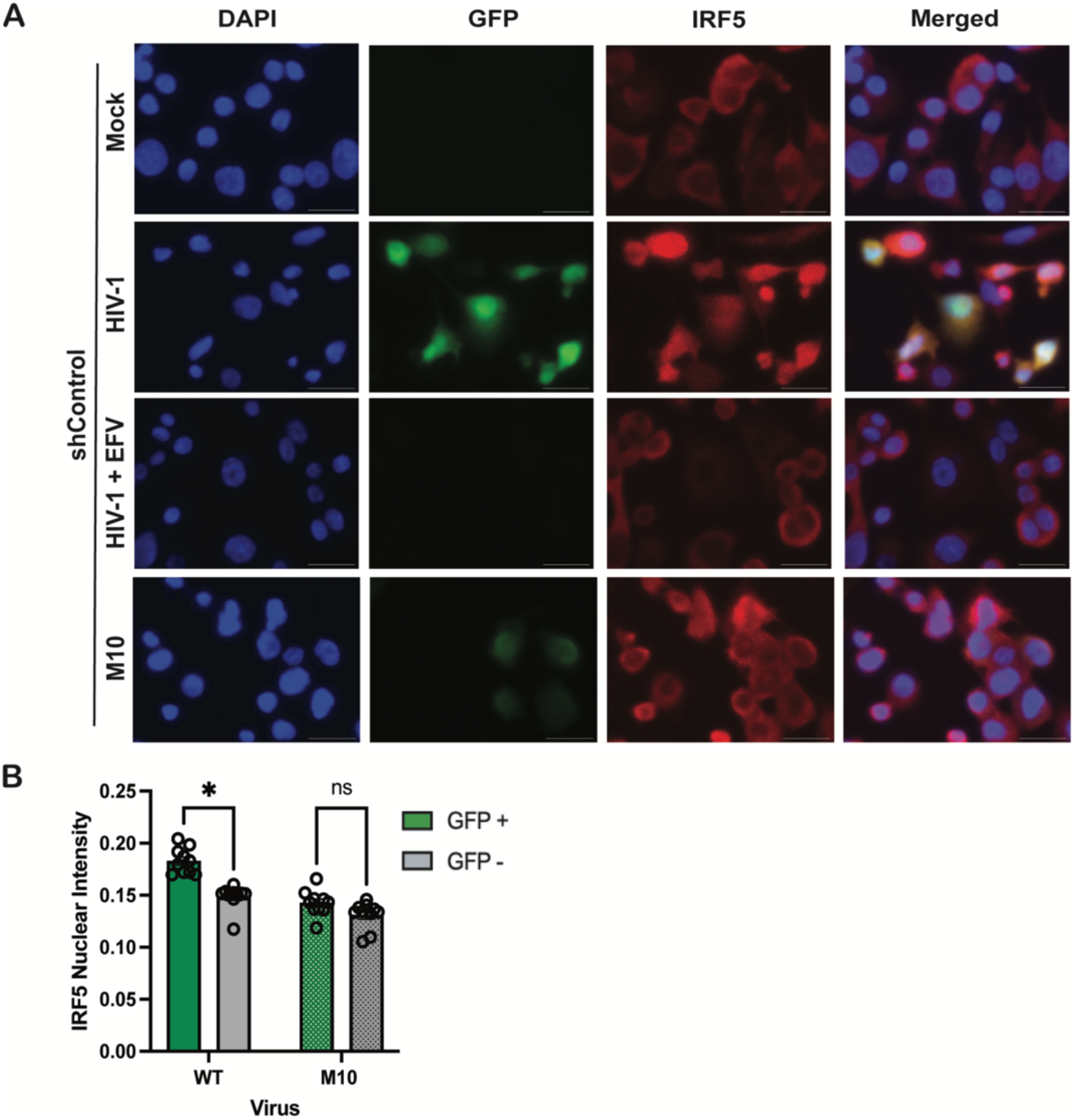
(A) THP-1/PMA macrophages were infected with LaiL’.envGFP/G (WT or M10 mutant) viruses (MOI 2). At 3 dpi, cells were fixed and stained for IRF5 localization and DAPI to visualize nuclei. Scale bar "’ 100 µm. (B) Cell Profiler was utilized to assess IRF5 nuclear intensity in infected and uninfected cells Images from three independent infection experiments were analyzed and quantified, with each dot representing a field containing approximately 50-150 cells. Statistical significance assessed via unpaired I-test (B). *: p < 0.05, ns ;a not significant.

**Supplementary Figure 5.**
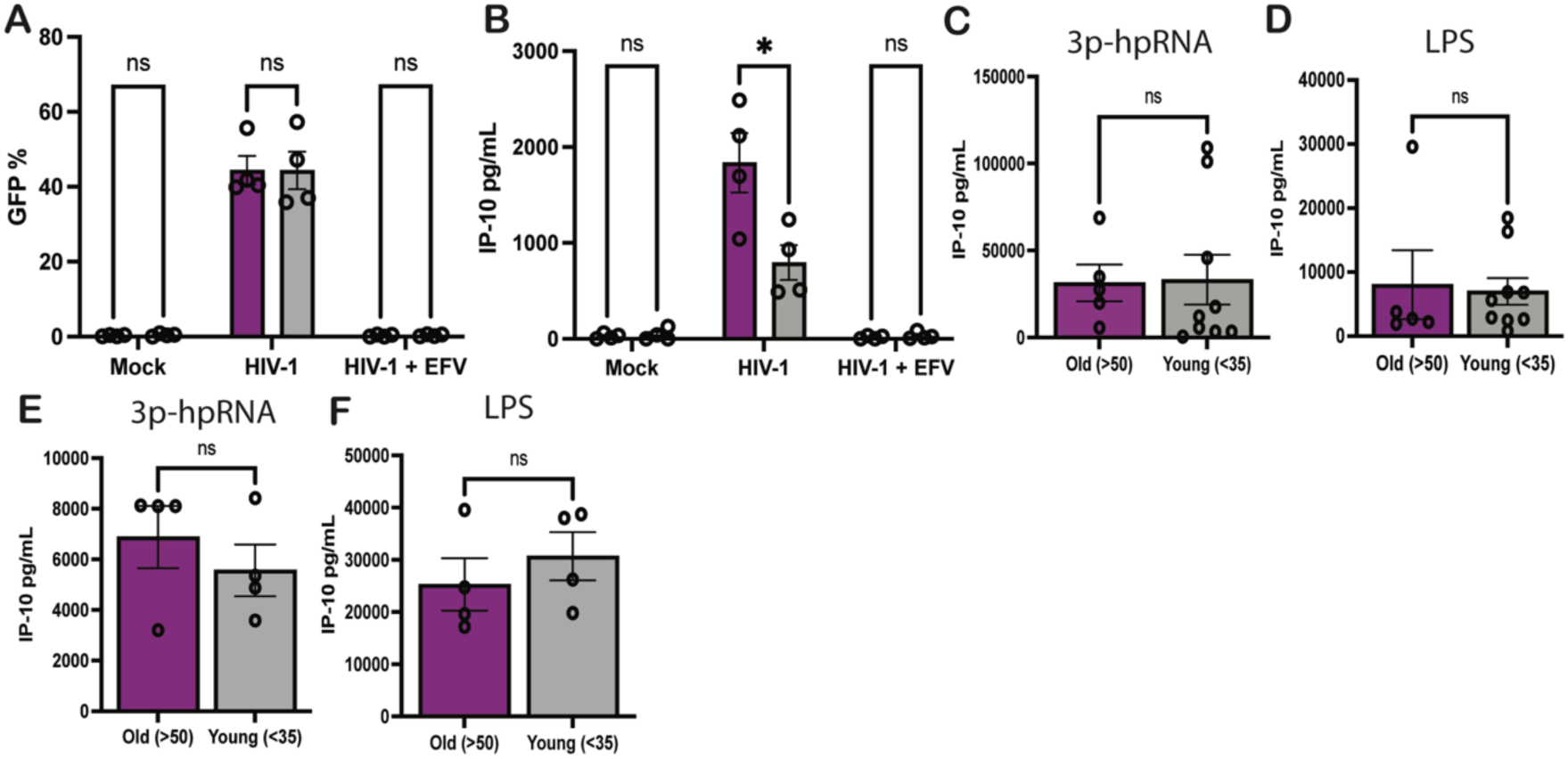
(A-8) MDMs (NY Biologics) were infected with Lai6.envGFP/G (MOI 1) in the presence of dNs and cells and supernatants were harvested at 3 dpi to assess (A) levels of infection and (8) IP-10 secretion. MDMs from HIV/Aging cohort (C-O) or NY Biologics (E-F) were treated with either 3p-hpRNA (2.5 ng/ml) or LPS (100 ng/ml) (G) and supernatant was harvested at 18 h post-stimulation to assess levels of IP-10 secretion by ELISA. Statistical significance was assessed via unpaired t-test (A-F) •: p<0.05, ns = not significant.

**Supplementary Figure 6.**
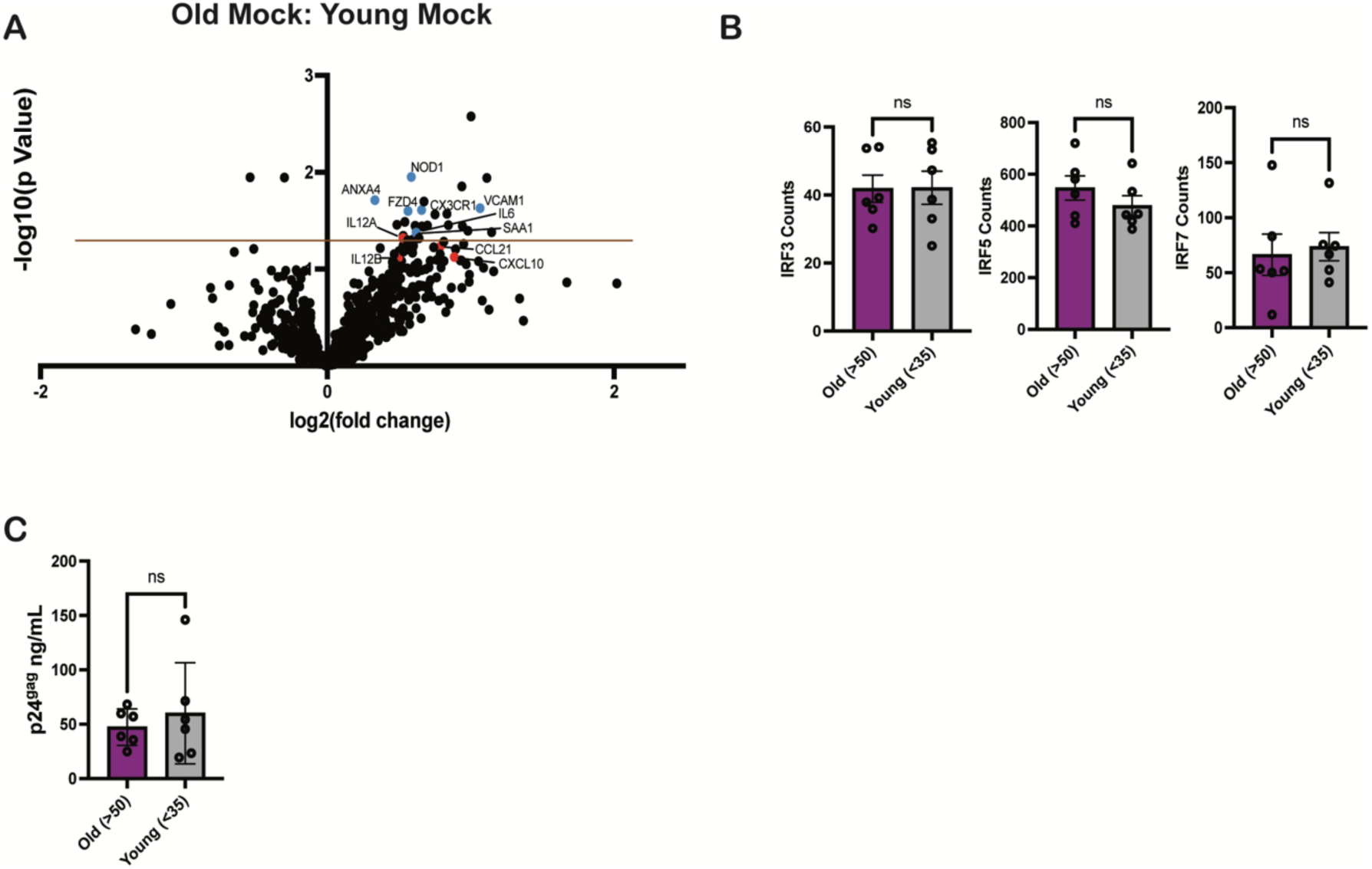
(A) RNA isolated from LaiMnvGFP/G-infected MDMs (MOI 2) was analyzed via Nanos­ tring nCounter using the Myeloid Innate Immunity V2 panel. Baseline expression of the gene panel was calculated using nSolver and plotted as a ratio of Old (Mock) vs. Young (Mock) with significantly upregulated IRFs (red) highlighted. The dashed line represents p-value of 0.05. (B) Raw count values for IRF3, IRFS, and IRF7 were plotted to assess differences in basal mRNA expression. (C) p24gag levels for selected donors in order to ensure equivalent levels of infection measured by ELISA Significance was assessed via unpaired two-tailed !-test (A-C). ns = not significant.

## Notes

### Competing Interest Statement

The authors have declared no competing interest.

### Summary of Updates

Order of the authors was incorrectly posted.

